# Human-like *APOBEC3* gene expression and anti-viral responses following replacement of mouse *Apobec3* with the 7-gene human *APOBEC3* locus

**DOI:** 10.1101/2024.07.30.605801

**Authors:** Nerissa K Kirkwood, Haydn M Prosser, Melvyn W Yap, Jane Gibson, Ross Cook, Ian Reddin, Ane Stranger, Emma Howes, Nur Zainal, Manikandan Periyasamy, Silvestro G Conticello, Gareth J. Thomas, James Scott, Kate N Bishop, Simak Ali, Allan Bradley, Tim R Fenton

**Affiliations:** School of Biosciences, University of Kent, Canterbury, Kent CT2 7NJ, UK; T-therapeutics. Abington Hall, Granta Hall, Granta Park, Cambridge, CB21 6AL; Retroviral Replication Laboratory, Francis Crick Institute, London, NW1 1AT, UK; School of Cancer Sciences, Faculty of Medicine, University of Southampton, Southampton SO16 6YD; Zoetis, Glenn Berge Building, Babraham Research Campus, Cambridge CB22 3FH; Department of Surgery & Cancer, Imperial College London, London W12 0NN, UK; Institute of Molecular and Cell Biology (IMCB), 61, Biopolis Dr, #06-12, Proteos, Singapore 138673; Core Research Laboratory, ISPRO-Institute for Cancer Research, Prevention and Clinical Network, 50139 Firenze, Italy; Institute for Life Sciences, University of Southampton, UK

**Keywords:** APOBEC3, mouse, cancer, virus

## Abstract

The seven human APOBEC3 (hA3) genes encode polynucleotide cytidine deaminases that play vital roles in restricting replication of viruses and retrotransposons. However, off-target A3 deamination of the cellular genome is a major source of somatic mutations in human cancer. The ability to study A3 biology *in vivo* is hindered by the fact that the solitary murine *Apobec3* gene (mA3) encodes a cytoplasmic enzyme, with no apparent mutagenic activity. Transgenic expression of individual hA3 genes in mice has helped to confirm their oncogenic potential but important questions including which hA3 genes are active in different tissue contexts and how they function in concert when under control of their cognate promoters cannot be addressed using these models. Here we describe humanization of the mouse mA3 locus by integration of a modified BAC clone encompassing the entire 7-gene hA3 locus from human chromosome 22 replacing mA3 on mouse chromosome 15. APOBEC3 mice are viable and fertile and hA3 gene expression in cells and tissues correlates strongly with expression in corresponding human cells and tissues, indicating human-like regulation of hA3 gene expression in the mice. Splenocytes from this line display a functional human A3 response to Friend Murine Leukaemia Virus (F-MLV) infection. We propose that the Hs-APOBEC3 mouse will uniquely model the function of the complete hA3 locus in a living organism and that it will serve as a useful background upon which to model human cancer, as well as assisting drug discovery efforts.

## Introduction

Humans possess seven Apolipoprotein B mRNA-editing enzyme catalytic polypeptide-like 3 (APOBEC3) genes (A3A, A3B, A3C, A3D, A3F, A3G, A3H), which reside in tandem in an ∼150 kb locus on chromosome 22 and encode zinc-dependent polynucleotide (deoxy)cytidine deaminases that catalyze the conversion of cytosine bases to uracil (C-to-U) in single-stranded DNA (ssDNA) and in some cases, RNA. Cells utilize both the mutagenic activities and non-enzymatic functions of these enzymes to inhibit viral and retrotransposon replication, making them an essential component of our innate immune system^1–3^.

Somatic mutation signatures in cancer genomes that are characterized by C>T and C>G substitutions at Tp<u>C</u> sites have been convincingly linked to off-target activities of A3A and A3B against ssDNA intermediates generated during processes including replication, DNA repair and transcription (reviewed in^4–8)^. Induction of mutagenic A3A and/or A3B activity in response to treatment has also been implicated in acquired resistance to cancer therapeutics^9–13^, while A3G, which deaminates cytosine in the Cp<u>C</u> dinucleotide context, has been implicated in generating a distinct mutational signature in cancer^14^. Although A3A and A3B appear to be responsible for the large majority of mutagenic A3 activity at Tp<u>C</u> sites, deletion of both enzymes from cancer cell lines does not eliminate the accumulation of A3 signature mutations in all cases, suggesting a contribution from one or more additional TpC-specific A3 enzymes^15^. A haplotype of A3H (A3H-I) has been proposed to contribute to cancer mutagenesis in certain contexts^16^ and a deaminase-dependent role in enabling pancreatic cancer cells to tolerate gemcitabine treatment by resolving stalled replication forks has also been described for A3C and A3D^17^.

Given its prominent role in mutagenesis, drugs that inhibit A3 activity could potentially be useful in cancer treatment, for example by suppressing the emergence of acquired drug resistance. However, the successful development of such a therapeutic strategy depends upon building a clear understanding of the isoform-specific roles and extent of functional redundancy between the A3 genes in physiologically relevant models of tumour development and treatment^18,19^. Studying A3 biology in mice has been hindered by the fact that whereas humans have seven A3 genes, mice have only one, *Apobec3* (mA3). mA3 inhibits murine retrovirus replication, however the mechanism appears to be cytidine deamination-independent^20–22^ and mA3 is a cytoplasmic protein^21,23^ as opposed to human A3A which can enter the nucleus and A3B, which is localized exclusively to the nucleus^24,25^. Furthermore, no mutational signature associated with mA3-mediated deamination of genomic DNA has been reported in mouse tumours. Expression of individual A3A or A3B transgenes under the control of constitutive or inducible heterologous promoters in mice has enabled confirmation of their oncogenic potential^26–29^ and their ability to promote acquired resistance to cancer therapeutics ^10^, and transgenic expression of A3A and A3G in mA3 knock-out mice has enabled comparison of their antiviral roles^30^. It is clear however, that A3 gene expression is normally tightly controlled and that enforced expression of either A3A or A3B is associated with significant cytotoxicity, both in cultured cells^25,31,32^ and in mice^10,26,33,34^, limiting the utility of such models for studying the physiological functions of these enzymes *in vivo*. The use of single A3 transgenes also necessitates a focus on one selected A3 enzyme in isolation from the other A3 paralogues, thus failing to recapitulate the complexity observed in humans which may be critical to understanding how best to suppress A3-driven mutagenesis in cancer patients.

Here we report the generation of a humanized A3 (Hs-APOBEC3) mouse line in which a modified BAC clone containing the entire hA3 gene locus was integrated by recombination-mediated cassette exchange into the mA3 locus on chromosome 15. The resulting mice possess all seven hA3 genes under the control of their own regulatory elements. To assess the utility of these mice for further studies we conducted RT-qPCR assays to measure the baseline expression levels of the hA3 genes in ES cell clones, selected tissues and peripheral blood mononuclear cells (PBMCs). Furthermore, the effect of known A3 inducers on the expression levels in ESCs and PBMCs was determined. Comparisons of these data with gene expression data from equivalent human cells and tissues showed strong correlations, indicating human-like regulation of hA3 gene expression in Hs-APOBEC3 mice. Both mA3 KO mice and Hs-APOBEC3 mice displayed a slightly shorter lifespan than WT littermates but no predisposition to spontaneous tumour development was apparent. Finally, mutagenesis of murine leukaemia virus (MLV) was observed upon infection of splenocytes from Hs-APOBEC3 mice, demonstrating a functional human A3 response to viral infection in this model. Taken together, our phenotypic characterization and analysis of hA3 gene expression and function in Hs-APOBEC3 mice suggest this line will serve as a valuable tool for the interrogation of A3 function in physiology and disease.

## Results

### Strategy for humanization of the mouse mA3 locus by recombination-mediated cassette exchange

The procedure used for humanizing the mA3 locus is outlined in Figure 1 and described in detail in *Methods*. To delete mA3 and create a locus suitable for insertion of human sequence a selection cassette flanked by loxP sites was targeted in mouse AB2.1 ES (129S7/SvEvBrd-Hprt^−^) cells^35,36^. As a 36kbp deletion was required to remove mA3, CRISPR constructs were designed to induce double stranded breaks in the chromosome^37^. Targeted ES cell clones lacking mA3 were microinjected into C57BL/6-*Tyrc-Brd* blastocysts to produce chimaeras that were then test-bred for germline transmission. Germline competent ES clones could then be used for transgenesis with the human BAC clone.

**Figure 1:**
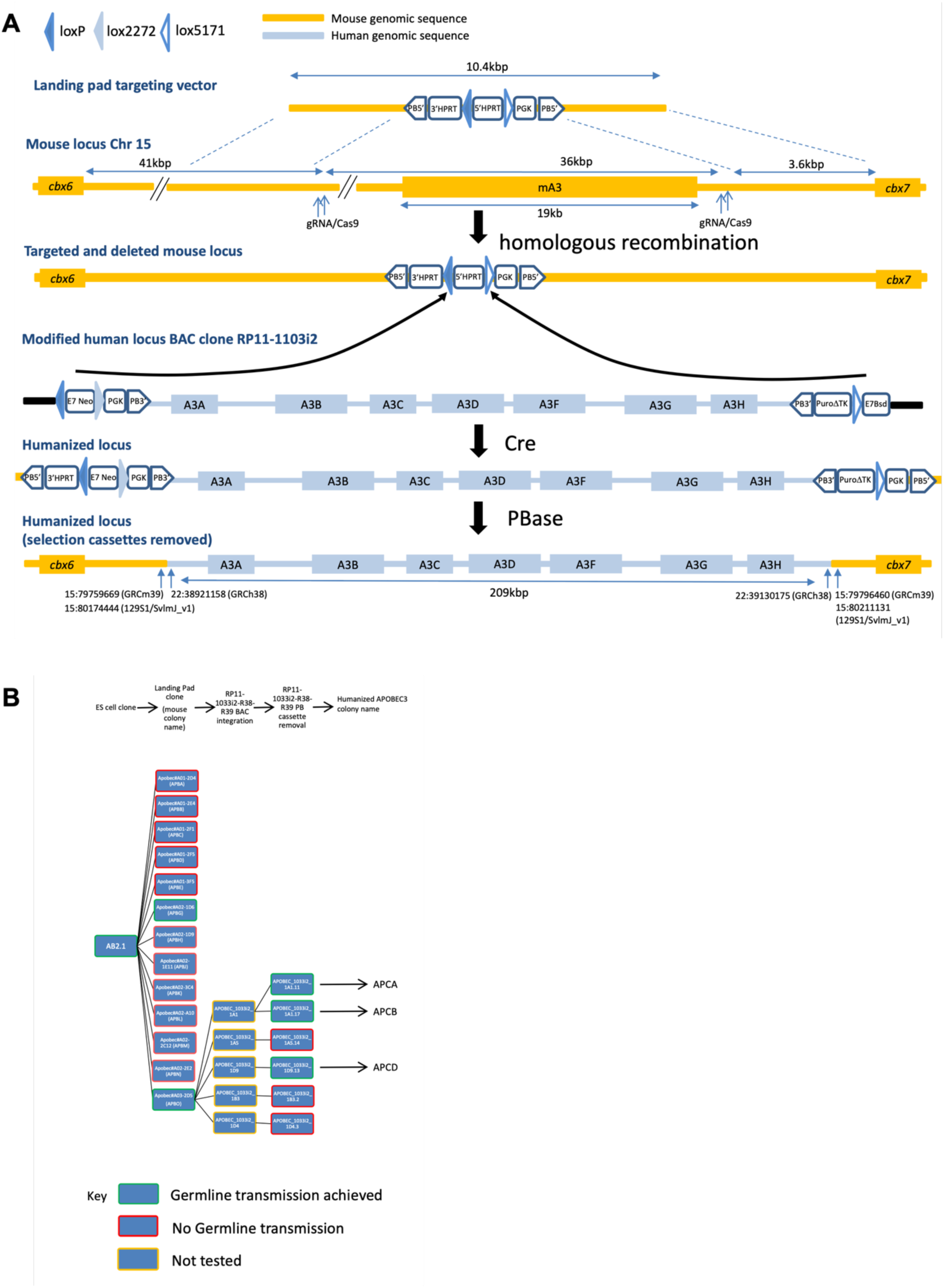
Schematic illustrating the steps used for humanizing the Apobec3 locus. (**A)** “Landing-pad” cassette was targeted by homologous recombination in mouse ES cells, deleting mouse APOBEC3 between 15: 79759669 to 79796460 (GRCm39) which is equivalent to 15:80174444 to 80211131 for 129S1/SvlmJ_v1. Human BAC clone RP11-1033i2 modified and trimmed by flanking cassettes to contain sequence between 22:38921158 to 39130175 (GRCh38), was co-transfected with a Cre expression plasmid insertion into the landing pad loxP and lox-5171 sites by RMCE was selected using Puromycin followed by G418. Finally the flanking cassettes were excised from the locus by transfection with a PBase expression plasmid and selection with FIAU. (**B)** Summary of relation between ES cell clones and mouse colonies for the sequential steps in humanizing the mouse APOBEC gene.

The human BAC clone RP11-1033i2 was modified by insertion of selection cassettes and recombinase sites at the flanks of the genomic insert by homologous recombination in *E. coli*^36,38^. Correct insertion of the recombineering cassettes and the integrity of the modified BAC clone was assessed by junction and internal PCR as well as Pulsed Field Gel Electrophoresis (PFGE) (Suppl. Fig. 1).

The modified RP11-1033i2 BAC was inserted into the landing pad as previously described^35,36^ by co-transfection with a Cre expression plasmid and ES cell colonies being selected with G418 followed by puromycin. Correct integration and integrity of the inserted BAC clone was assessed by junctional and internal PCR (Suppl. Fig. 2). ES cell clones were transfected with a piggyBac transposase (PBase) expression plasmid^39,40^ to effect deletion of both recombineering cassettes, loss of the 3’ flanking cassette being selectable for loss of the PuTK gene with 2′-fluoro-2′-deoxy-β-D-arabinofuranosyl-5-iodouracil (FIAU). Excision of recombineering cassettes was assessed by PCR on a clonal basis. Heterozygous APOBEC3 humanized ES cell clones were used to make chimaeras, and upon germline transmission, the humanized allele was bred to homozygosity.

### Hs-APOBEC mouse ESCs express human A3 genes

As a first step in assessing the utility of the humanized A3 mice, we mapped human A3 gene expression in the mESCs used in their generation. This included ES clones APOBEC_1033i2_1A1_11 (1A1_11), APOBEC_1033i2_1A1_17 (1A1_17) and APOBEC_1033i2_1D9_13 (1D9_13). Furthermore, we looked at hA3 expression in ES clone APOBEC_1033i2_1A4_23 (1A4_23), which was not used for the subsequent mouse generation. All seven human A3 genes were expressed in the mESCs to varying degrees, with A3B and A3C showing the highest levels of expression and A3A, A3D and A3F the lowest (Figure 2A, 1A1_11; Suppl. Figure 3A-C, 1A1_17, 1A4_23, 1D9_13). Comparison of hA3 gene expression in the mESCs with RNA-seq data from hESC line H1 (GEO accession GSE75297) revealed consistent expression patterns for most hA3 genes between the two cell types, except for A3D and A3F which were expressed at lower levels in the mESCs (Figure 2B, 1A1_11; Suppl. Figure 3D-F, 1A1_17, 1A4_23, 1D9_13). Pearson’s *R*-values indicate strong positive correlations between hA3 gene expression in the transgenic mESC clones and hESC line H1 (0.65, 1A1_11; 0.73, 1A1_17; 0.65, 1A4_23; 0.74, 1D9_13). These values increase to between 0.82 and 0.88 upon removal of A3D and A3F from the analysis (Suppl. Figure 3G-J).

**Figure 2.**
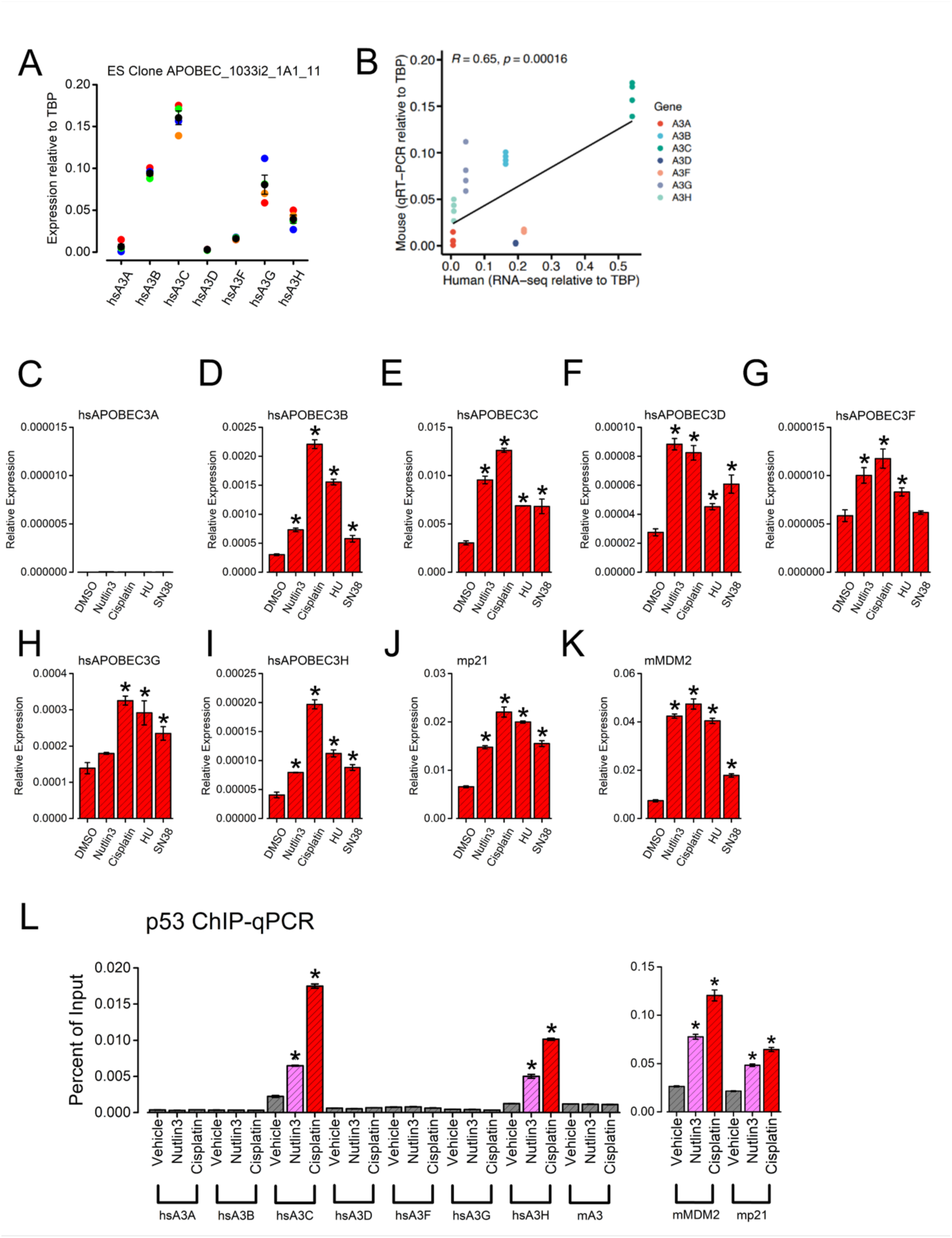
Correspondence between hA3 gene expression in human and Hs-APOBEC3 mouse ES cell clones. (**A**) Baseline hA3 gene expression levels in ES cell clone 1A1_11. Expression levels are normalized to the reference gene *Tbp* (n=4; red, repeat 1; orange, repeat 2; blue, repeat 3; green, repeat four; black, mean). (**B**) hA3 gene expression in mouse ES cell clone 1A1_11 compared with RNA-seq data from human ES cell clone H1. (**C-K**) Graphs showing the effects of Nutlin-3A (10 µM), cisplatin (20 µM), hydroxyurea (HU, 2 mM) or SN-38 (50 nM) on hA3 (**C-I**), mp21 (**J**) and mMDM2 (**K**) gene expression levels in mouse ES cell clone 1A4_23. RT-qPCR data is shown relative to GAPDH levels (n=3). Significantly (*p*<0.05, one-way ANOVA) higher gene expression levels, relative to the vehicle control, are indicated (*). (**L**) p53 ChIP following addition of Nutlin3 or cisplatin for 24 hours to mouse ES cell clone 1D9-13 (n=3). Significant changes (*p*<0.05) relative to the vehicle control are indicated (*). Error bars show SEM.

### Chemotherapeutic drugs induce human A3 gene expression in Hs-APOBEC3 mouse ESCs

Cytotoxic drugs that interfere with DNA replication promote expression of several A3 genes, most notably A3B, A3C and A3H, in human breast and colon cancer cell lines^12,41^. To determine if hA3 expression is controlled in a similar manner in our Hs-APOBEC3 mESCs, hA3 expression levels were measured in cultured cells (clone 1A4_23) following 24-hour exposure to either the vehicle control (DMSO) or to Nutlin-3A (which stabilizes p53 by inhibiting its MDM2-mediated ubiquitination), the DNA crosslinker, cisplatin, the ribonucleotide reductase inhibitor, hydroxyurea (HU) or the topoisomerase inhibitor, SN-38. Except for A3A, which is not expressed in either mESCs or in hESCs (Figure 2B), all drug treatments stimulated expression of one or more A3 genes (Figure 2C-I). Given their established ability to induce p53, all treatments also promoted the expression of p53 target genes, *Cdkn1a* (mp21, Figure 2J) and *Mdm2* (Figure 2K).

A3C and A3H are p53 target genes ^42,43^ and both Nutlin-3A and cisplatin promote p53 recruitment to the A3C and A3H promoters in human cancer cell lines^12^. Chromatin immunoprecipitation from Nutlin-3A or cisplatin-treated hA3 mESCs (clone 1D9-13) also revealed recruitment of mouse p53 to the A3C and A3H gene promoters (Figure 2L, left). Taken together with the qRT-PCR data (Figure 2F, J), these experiments demonstrate that regulation of A3C and A3H by p53 is conserved in our hA3 mouse model. As expected, p53 was also recruited to the *Cdkn1a* (mp21) and *Mdm2* promoters following Nutlin or cisplatin treatment (Figure 2L, right).

### Hs-APOBEC3 mouse tissues display human-like expression of the hA3 genes

We next mapped basal hA3 gene expression in selected tissues (ovary, lung, spleen) isolated from the homozygous humanized mice. Mice from all three lines were included (APCA, APCB, APCD). We measured expression of all hA3 genes in each tissue, with A3C consistently showing the highest levels and A3D the lowest (Figure 3A-C, APCA ovary, lung, spleen; Suppl. Figure 4A, APCB ovary, lung, spleen; Suppl. Figure 4B, APCD ovary, lung, spleen). Comparisons of the hA3 gene expression levels between the humanized mice and the corresponding human tissues using data from the Genotype-Tissue Expression (GTEx) Portal are shown in Figure 3D-F (APCA ovary, lung, spleen), Suppl. Figure 4C (APCB ovary, lung, spleen) and Suppl. Figure 4D (APCD ovary, lung, spleen). Pearson’s *R*-values range from 0.83-0.95 (ovary), 0.8-0.85 (lung) and 0.59-0.68 (spleen) revealing strong similarity between basal hA3 gene expression patterns in these tissues between the Hs-APOBEC3 mice and humans.

**Figure 3.**
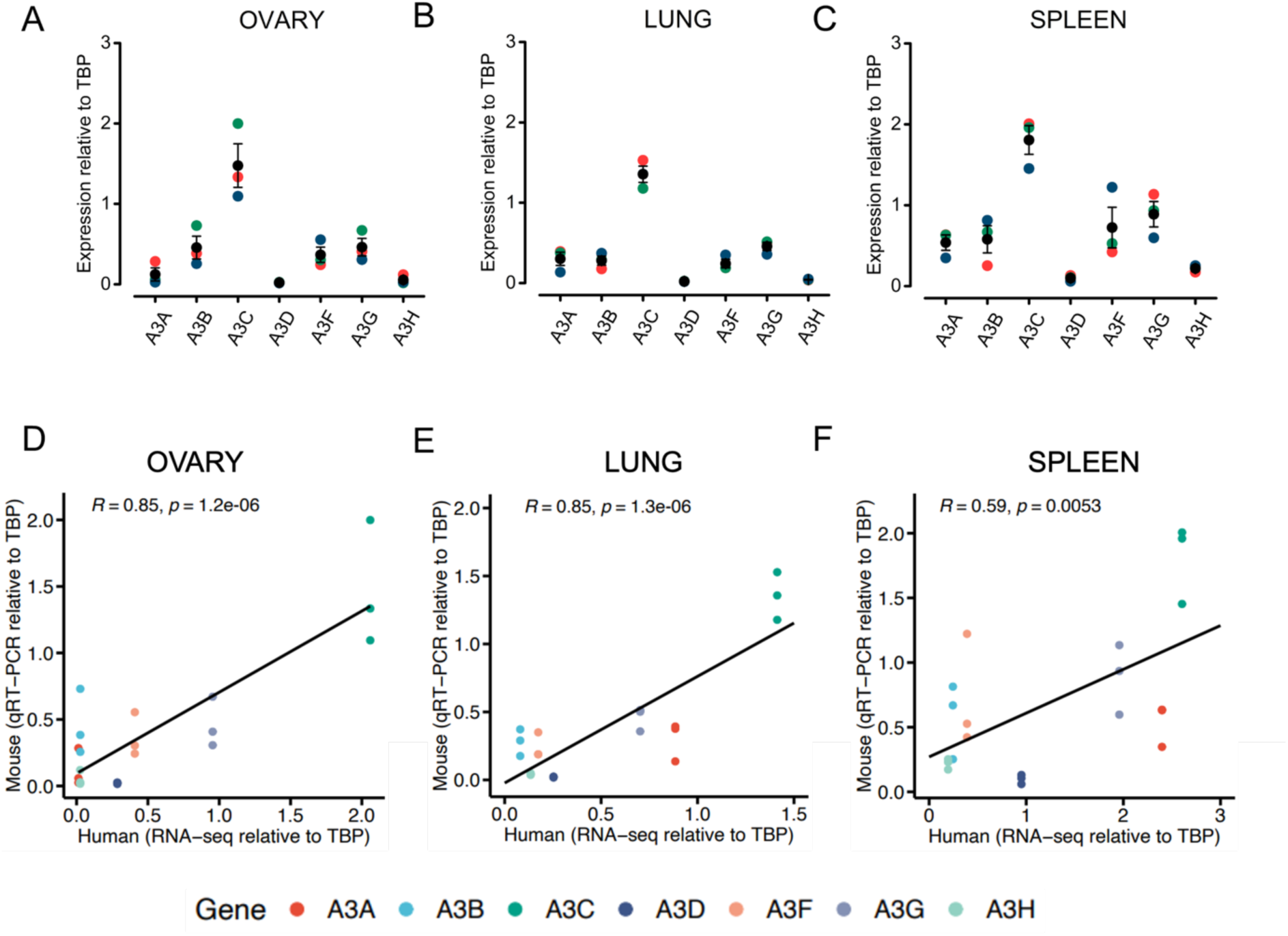
Human and humanized mouse tissues express hA3 genes at comparable levels. (**A-C**) Baseline hA3 gene expression levels in the ovary (**A**), lung (**B**) and spleen (**C**) of adult homozygous APCA mice. RT-qPCR data are shown relative to *Tbp* (n=3; mouse 1, red; mouse 2, orange; mouse 3, green; mean, black). Error bars show SEM. (**D-F**) hA3 gene expression in the humanized mouse tissue compared with the corresponding human data from the Genotype-Tissue Expression (GTEx) Portal for the ovary (**D**), lung (**E**) and spleen (**F**). Pearson correlation coefficients are indicated for each tissue.

### Human-like A3 gene expression and IFN induction in PBMCs from Hs-APOBEC3 mice

PBMCs are a heterogeneous cell population consisting of lymphocytes, monocytes and dendritic cells. We isolated PBMCs from the homozygous Hs-APOBEC3 mice and consistent with previous measurements of A3 gene expression in human PBMCs^44^, we observed high A3A and low A3B levels in unstimulated cells (Figure 4A). Comparison of our expression data from Hs-APOBEC3 PBMCs with RNA-seq data derived from human PBMCs^45^ revealed a strong correlation between hA3 gene expression between the two datasets (Figure 4B, Pearson’s *R*=0.89, p=6.8E-15). Subsequently, we isolated PBMCs from Hs-APOBEC3 mice to test the induction of hA3 genes *ex vivo* following treatment with known A3 inducers including the phorbol ester, phorbol 12-myristate 13-acetate (PMA) and interferon alpha (IFNα, Figure 4C, Suppl. Figure 5). Most notably, murine IFNα (mIFNα) induced over 5-fold increase in A3A expression (Figure 4C, Suppl Figure 5A), a finding in agreement with previous reports detailing the effect of IFNα on A3A expression in human PBMCs^44,46–50^. Human IFNα, which does not bind to the mouse type 1 IFN receptor^51^ had no effect on hA3 expression (Figure 4C, Suppl. Figure 5A-G). PMA’s effect on hA3 gene expression appears to be cell-type dependent; it potently induces A3A in human keratinocytes^52–54^ but in the MCF10A cell line derived from mammary luminal epithelium, it was found to induce A3B but not A3A^55^. A combination of PMA and IL-2 was found to decrease deaminase activity attributable to A3A in human PBMC extracts^50^ and consistent with this, we saw a decrease in A3A expression upon treatment of PMBCs from Hs-APOBEC3 mice with PMA (Figure 4C, Suppl. Figure 5A).

**Figure 4.**
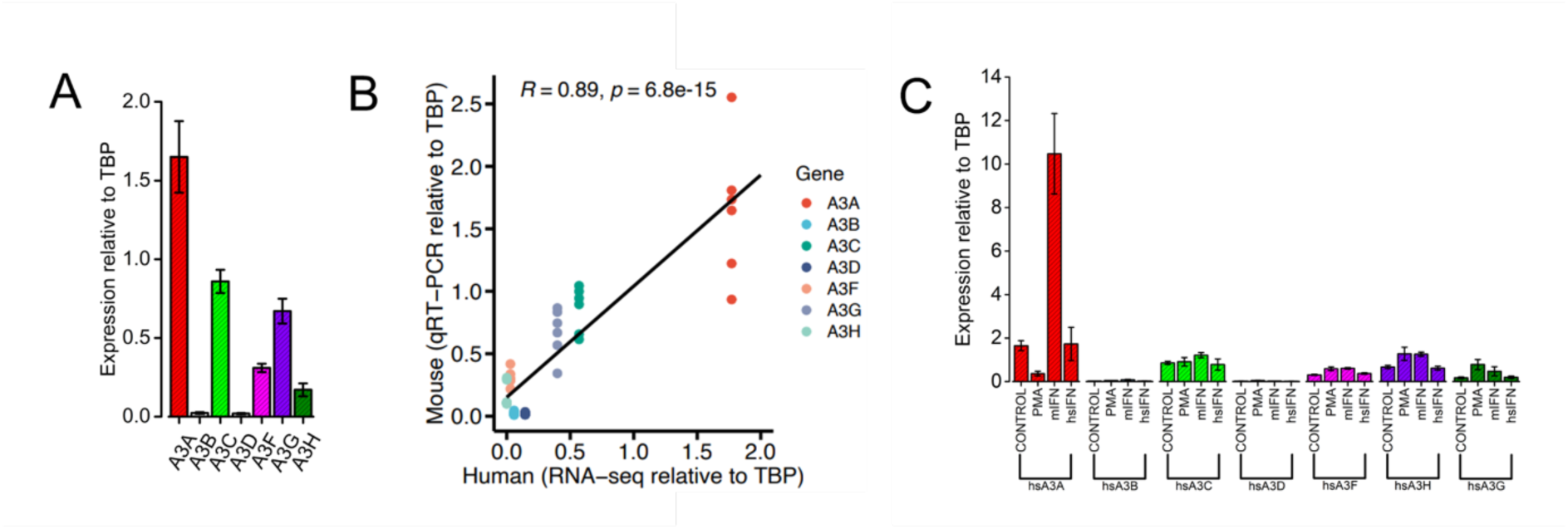
hA3 gene expression is inducible in PBMCs from the humanized mice. **(A)** Baseline hA3 gene expression levels in PBMCs isolated from adult homozygous APCA mice. RT-qPCR data is shown relative to *Tbp* (n=6). **(B)** A comparison between the hA3 expression levels in the PBMCs from the humanized mice and RNA-seq data from PBMCs isolated from four healthy volunteers reveals a strong correlation between the two (*R*=0.89). **(C)** hA3 gene expression levels in the humanized mouse PBMCs following 6-hour exposure to control media (n=6), PMA (n=4), mouse IFNα (n=3) or human IFNα (n=3) shown for all 7 hA3 genes. Error bars show SEM.

### *Apobec3* KO and *Hs-APOBEC3* mice display decreased *Cbx7* expression but lack phenotypic changes previously associated with *Cbx7* loss

In both mouse and human, the A3 locus is flanked by genes encoding the polycomb-group (PcG) proteins, Cbx6 and Cbx7. To check for any disruption to these genes following the deletion of mouse A3 and the subsequent integration of the human A3 locus, expression levels of *Cbx6* and *Cbx7* were compared in ovary, lung and spleen from all genotypes (Figure 5A-F). *Cbx7* showed significantly reduced expression levels in all tissues from mice of all genotypes compared to the WT mice, apart from spleens from the HET mice (Figure 5D-F). *Cbx6* displayed significantly reduced expression in all tissues from the HOM mice, in the ovaries and lungs of the HET mice and only in the lungs of the mA3 KO mice (Figure 5A-C). *Cbx6* expression was significantly increased in spleens from the mA3 KO mice compared to the WT mice (Figure 5C).

**Figure 5.**
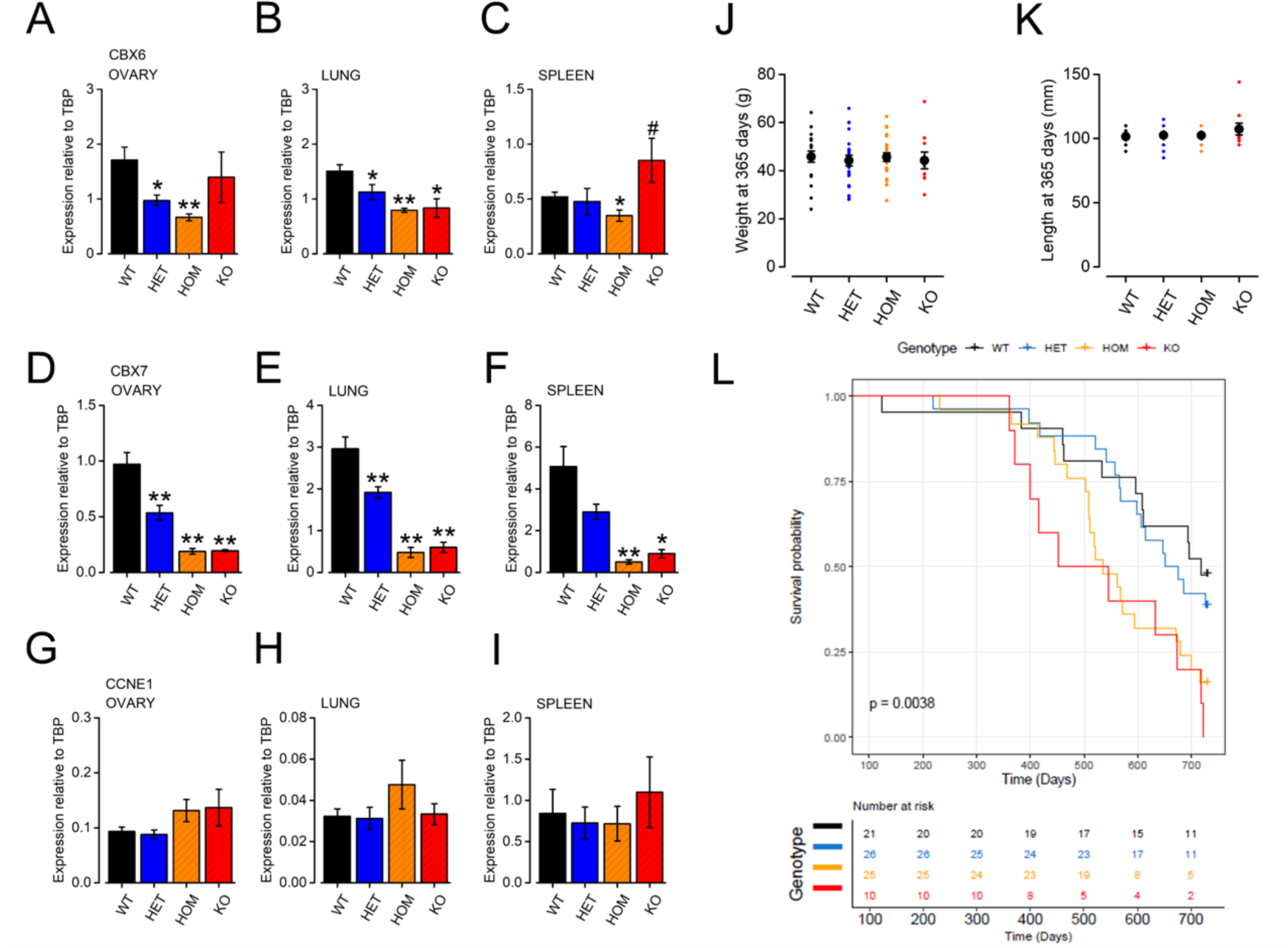
Hs-APOBEC mice do not display a *Cbx7* KO phenotype. (**A**) Kaplan-Meier survival curves for WT (n=21), heterozygous (n=26), homozygous (n=25) and mA3 KO (n=10) mice aged to a maximum of 2 years. (**B-G**) Expression levels of the two genes flanking the mA3 locus, *Cbx6* and *Cbx7*, in the ovary (**B, E**), lung (**C,F**) and spleen (**D,G**) of WT (n=9), heterozygous (n=9), homozygous (n=9; with the exception of homozygous spleen, n=8) and mA3 KO (n=3) mice. Expression levels are normalized to *Tbp*. Significantly lower (*p*<0.05, *; *p*<0.01, **) or higher (*p*<0.05, #) expression levels, relative to WT, are indicated. (**H-J**) *Ccne1* expression levels, normalized to *Tbp*, in the ovary (**H**), lung (**I**) and spleen (**J**) of the same mice. (**K, L**) Weight (g) and length (mm) measured at 365 days of mice on the aging study from all genotypes. Error bars show SEM.

A previous study of *Cbx7*-KO mice revealed overexpression of *Ccne1*, a gene encoding the protein cyclin E1, increased naso-anal length (measured at 12 months) and a pre-disposition to liver and lung adenomas and carcinomas in aged animals (17-23 months)^56^. To check for a *Cbx7* phenotype in the KO and humanized mice, we compared *Ccne1* expression levels in tissues (ovary, lung, spleen) from all genotypes. There was no significant difference in *Ccne1* expression in any tissue from any genotype compared to the WT levels (Figure 5G-I). Patterns of *Cbx6*, *Cbx7* and *Ccne1* expression in ovary, lung and spleen were consistent across all three Hs-APOBEC3 lines (APCA, APCB and APCD, Suppl. Figure 6). Comparison of the weight and length of all mice measured at 365 days of age revealed no differences between genotypes (Figure 5J, K).

To evaluate longevity and to check for any predisposition to tumour development as reported for *Cbx7*-KO mice^56^, we entered a cohort of 82 mice from all genotypes into an aging study up to a maximum of two years (number of mice: WT 21; HET 26; HOM 25; KO 10). Kaplan-Meier survival curves show the different survival times for each of the four genotypes (Figure 5L). Hazard ratios (HR) calculated for each genotype relative to the WT mice reveal a reduction in life expectancy for mA3 KO and homozygous hA3 mice with the KO mice having the greatest probability of death over the study period (HR values: homozygous hA3 = 2.56 (95% CI 1.23 - 5.35, p = 0.012); mA3 KO = 3.47 (95% CI 1.46 – 8.22, p = 0.005). CBX7 has been implicated in extending cellular lifespan by downregulating *INK4/ARF* expression and therefore inhibiting replicative senescence^57^, so it is possible that a loss of this activity in mA3 KO and homozygous hA3 mice could explain their reduced longevity relative to WT littermates.

Autopsy of end-stage animals revealed benign liver adenomas in 13 of 82 mice (4/25 WT, 6/25 HET, 2/25 HOM and 1/10 KO) but no obvious malignancies. We conclude that replacement of mA3 with the hA3 locus does not predispose to tumour development, either via direct effects of hA3 expression or indirectly, through reduced *Cbx7* expression.

### Evidence for functional human A3 responses to F-MLV in splenocytes from Hs-APOBEC3 mice

The mA3 enzyme of C57BL/6 mice restricts F-MLV replication via a deaminase-independent mechanism and does not generate G>A mutations in the MLV genome ^58–61^. To test whether one or more hA3 genes could respond to MLV in Hs-APOBEC3 mice, splenocytes were isolated from WT, heterozygous or homozygous Hs-APOBEC3 animals, stimulated with lipopolysaccharide (LPS) and infected with Friend MLV (F-MLV). Infected splenocytes were then co-cultured with *Mus dunni* tail fibroblasts (MDTF) to allow detection of viral genomes by qPCR after two or five days. Genomic DNA was also extracted after five days of co-culture and sequenced for mutations (Figure 6A). After two days of co-culture, viral titres as measured by qPCR were lower in *Mus dunni* cells co-cultured with homozygous Hs-APOBEC3 mouse splenocytes than WT or heterozygous splenocytes (one-way ANOVA, p=0.0653). However, by 5 days of co-culture, there was no difference between samples (Figure 6B, C). This indicates that either residual unrestricted virus was able to grow out in all mice over time, or that equivalent F-MLV restriction occurs in both WT and homozygous Hs-APOBEC3 mice. Given that deletion of mA3 results in increased F-MLV infectivity in C57BL/6 mice ^59^, our results suggest that one or more of the hA3 genes might be able to compensate for the defect in F-MLV restriction associated with mA3 loss. Long-read sequencing of a 2,186bp amplicon spanning the *env* gene at the 3’ end of F-MLV (a hotspot for APOBEC3G mutagenesis in the closely related Moloney MLV and other retroviruses^62^) revealed significantly more G>A mutations in viral genomes following infection of splenocytes from homozygous Hs-APOBEC3 mice (Figure 6D; Pearson’s chi squared test, p=3.55−10^−7^). G>A mutations observed in viral genomes isolated from Hs-APOBEC3 splenocytes were enriched in the <u>G</u>pG dinucleotide sequence context (Figure 6E; Pearson’s chi squared test, p=0.008), which is consistent with deamination of Cp<u>C</u> sites by APOBEC3G. When comparing the number of G>A mutations per sequencing read (i.e. mutations on the same cDNA molecule), we saw enrichment in F-MLV genomes from Hs-APOBEC3 homozygous and heterozygous splenocytes at <u>G</u>pG (Figure 6F, Kruskal-Wallis test WT vs HOM, p<2.2−10^−16^;; WT vs HET, p=4.7−10^−5^) and to a lesser extent, also at <u>G</u>pA dinucleotides (Kruskal-Wallis test WT vs HOM, p<2.2−10^−16^; WT vs HET, p=3.9−10^−7^); the latter implicating one or more of the Tp<u>C</u>-specific hA3 enzymes in responding to F-MLV. This is reminiscent of the response to HIV-1 observed in human cells, in which A3G and A3F generate G>A mutations through deamination of CpC and TpC sites respectively ^63,64^.

**Figure 6.**
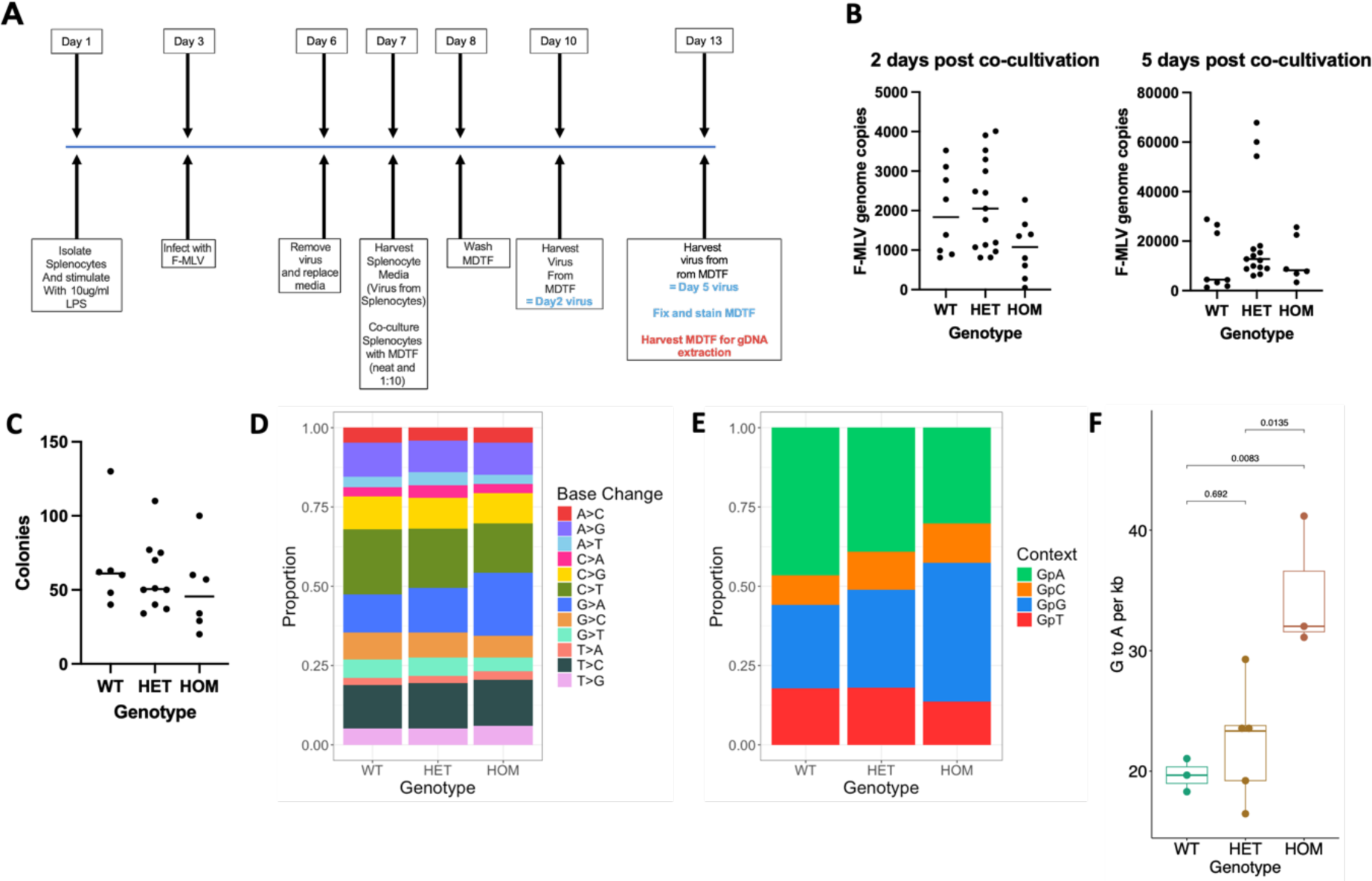
Functional responses to F-MLV infection in splenocytes from Hs-APOBEC3 mice. **(A)** Schematic representation of the experimental workflow employed for measuring F-MLV infectivity and APOBEC responses in splenocytes. (**B**) qPCR-based measurement of F-MLV titres in supernatant collected from *M. dunni* tail fibroblasts (MDTF) at 2 days (top) and 5-days (bottom) post-co-cultivation with infected splenocytes. (**C**) Colony counts from MDTF cultures at 5-days post-co-cultivation with infected splenocytes. (**D**) Stacked bars represent the proportion of different SNVs detected in a 2,186 bp amplicon from the F-MLV *env* gene following infection of splenocytes from WT, HET or HOM mice. There is a significant difference (Pearson’s chi squared test, p=3.55−10^−7^) in G>A (GA, blue) mutations between genotypes, which analysis of residuals indicates is driven by an increase in G>A in the HOM group. (**E**) Stacked bars represent the proportion of G>A mutations observed in different dinucleotide sequence contexts. There is a significant difference (Pearson’s chi squared test, p=0.008) between groups, which analysis of residuals indicates is driven by an increase in G>A mutations at GpG sites (blue) in the HOM group. (**F**) Comparison of G>A mutation rates per kilobase in F-MLV *env*. P-values for pairwise comparisons calculated using one-way ANOVA with Tukey’s post-hoc test.

## Discussion

Several A3 transgenic mice have been reported in which expression of a single human A3 gene is driven by a heterologous promoter, either on a wild-type ^29,33,34^ or *Apobec3* knockout ^26,30^ background. These mice have been useful for studying aspects of A3 biology (e.g. the oncogenic potential of A3A and A3B and their ability to generate murine tumours displaying the APOBEC mutation signatures ^26,29^, or the viral restriction activities of A3A and A3G ^30^). However, they only enable the study of one A3 gene in isolation from the remaining paralogues, nor do they recapitulate the normal regulation of A3 gene expression, due to the use of heterologous promoters for driving transgene expression. In our model, we demonstrate the co-expression of multiple A3 genes and observe regulation of expression that is similar to that seen in human cells and tissues. Preservation of physiological A3 gene regulation is likely to be particularly important when attempting to generate mouse tumours that harbour high APOBEC-induced somatic mutation burdens and that are therefore more antigenic (and in turn, suitable for studying cancer immunology and immunotherapy) than conventional mouse tumour models. In the case of the A3A and A3B transgenic mice that have been utilized thus far, the toxicity associated with expression of either gene from heterologous promoters has limited researchers to the use of models in which expression is sufficiently low to avoid acute toxicity to the animal (in the case of conditional expression in adult mice ^33^) or to avoid selection against cells expressing the functional transgene during development ^26,30^. This low level of constitutive expression is quite different to the considerable fluctuations in A3A and A3B expression that are observed during cell cycle, cellular differentiation, viral infection and interferon signalling or in response to cancer therapeutics ^10–12,41,65–70^, and which likely underlie the generation of APOBEC mutation signatures *in vivo*.

There are some important limitations to note in this Hs-APOBEC3 model. In the interests of recapitulating A3 gene regulation and function as faithfully as possible, we incorporated the unmodified human locus, thus our model lacks fluorescent reporters or epitope tags that could be useful in monitoring A3 expression or activity. These are modifications that could be engineered into the line, together with targeted disruption of individual A3 genes, or generation of the *APOBEC3A_B* deletion allele that has been associated with altered cancer risk and to increased somatic mutagenesis in breast cancer, potentially due to increased A3A and/or A3H activity. Finally, although we have demonstrated human-like A3 expression patterns in selected cells and tissues, much remains to be learned about A3 regulation in human cells and it is unclear to what extent all facets of regulation (both of gene expression and protein function) will be mirrored in the Hs-APOBEC3 mouse. Nonetheless, the data presented here suggests the Hs-APOBEC3 line will be a useful tool for studying A3 biology *in vivo* and that it may serve as a background on which human cancer and cancer treatment can be more accurately modelled than in existing wild-type or single A3 transgenic mice.

## Methods

### Ethics

All animal breeding and procedures were conducted following local ethics approval, under the following UK Home Office Project Licences: PD165A873 (generation of Apobec3 KO and Hs-APOBEC3 mice and establishment of breeding colonies); PA0478C6F and PP3714531 (breeding and collection of tissues); PP1364481 (ageing study).

### Gene targeting of mA3

A targeting vector was created to facilitate deletion of mA3 and insertion of a cassette with recombinase sites, or “landing pad” that can integrate the human locus (Fig. 1a and Suppl Figure 1A). The targeting vector was constructed by PCR of a 2988bp 5’ targeting arm from BAC RP23-373B5 (using primers mAPOBEC5.1 and mAPOBEC5.2). A 129 derived BAC was not used for this amplification due to what was later found to be a 3134bp deletion in 129 genome relative to the reference C57Bl/6 genome which deleted the 5’ end of the 5’ targeting arm. A 2957bp 3’ targeting arm was amplified from BAC bMQ-365P11 ^71^ (using primers mAPOBEC3.1 and mAPOBEC3.2). A four-piece Gibson reaction (NEB) was performed using the two targeting arms, a cassette excised from a modified pRMCE23 vector ^36^ and a pBluescript (Agilent) based vector backbone to create pAPOBEC3_TV. pAPOBEC3_TV was linearized with PmeI (NEB), ethanol precipitated and re-suspended in TE in preparation for transfection. As the targeting vector was designed to delete 36,759bp of mouse sequence, efficiency was to be enhanced by using CRISPR. Four double stranded oligo nucleotides were cloned into the pX330 vector ^37^ at the BbsI site to create gRNAs for site specific cutting. Two gRNAs were designed to target each end of the mouse *Apobec3* deletion. The gRNA vectors were used in pairs, one targeting each end, and in four different combinations.

The four DNA mixes were electroporated into AB2.1 ES cells using (program A23) AMAXA (Lonza). In each case 1μg of each of the two gRNA vectors and 10μg of pAPOBEC3_TV was used with 5 x 10^6^ ES cells. HAT selection was applied after 24 hours and continued for a further 8 days, being replaced at that point by HT supplement. In total 576 surviving colonies were picked into 96-well plates, expanded into three copies two being aneuploidy for chromosomes frozen and a third being digested with proteinase K and the genomic DNA being precipitated, dried and re-suspended in TE for genotyping (Suppl. Figure 1B). As it was difficult to identify genotyping primers without non-specific amplification, the genomic DNA was initially digested with XmnI and transferred by Southern blot to a nylon membrane (Thermo) for hybridization with a 3’ external probe. Clones that had hybridizing XmnI fragments indicative of heterozygous targeting of the mA3 locus were then PCR amplified by primers spanning the junction between the cassette and the mouse genomic sequence outside of the targeting arms at the 5’ and 3’ ends. Clones passing all three genotyping stages were then checked by qPCR for absence of Y, 8 and 11. Individual ES cell clones were then micro-injected into blastocysts to create chimaeric mice to assess germ line competence. Heterozygote mice were bred with each other to generate homozygotes which were checked for deletion of Apobec3 exons by PCR amplification (Suppl. Figure 2).

### BAC transgene preparation for humanization

In preparation for introducing the hA3 locus a human BAC clone RP-11-1033i2 ^72^ spanning the locus was obtained. To trim extraneous DNA from the ends of the recombineering vectors were constructed based upon pRMCE38 and pRMCE39 ^36^. pRMCE 38 was modified by replacement of the 3’ piggyBAC (PB) LTR with a 5’ PB LTR. As the selection cassettes for both pRMCE38 and pRMCE39 will now be flanked by PB 5’ and 3’ LTRs this will enable removal of the recombineering cassette by PBase ^39,40^. Recombineering arms were then PCR amplified from the RP11-1033i2 BAC and cloned into the pRMCE38 and pRMCE39 vectors by ligation. The recombineering enabling vector pSim18 ^38^ was transformed into RP11-1033i2 containing *E. coli*, followed by the excised and gel eluted recombineering cassettes from the modified pRMCE38 and pRMCE39 vectors (Suppl. Figure 7). The trimmed BAC with the cassettes inserted (RP11-1033i2-R38-R39) were checked by PCR across the recombineering junctions followed by restriction digest and pulse-field gel electrophoresis. RP11-1033i2-R38-R39 was prepared for transfection using Large Costruct Kit (Macherey-Nagel).

### BAC transgenesis for APOBEC3 humanization

One germline competent landing pad clone Apobec#A03-2D5 was transfected (Bioroad) with 10μg of BAC DNA and 20μg of pCAG-iCre ^73^. Colonies that had integrated the modified RP11-1033i2 BAC were selected with G418 (180μg/ml) for 5 days, followed by Puromycin (3μg/ml) for another 5 days. Surviving colonies were picked into 96-well plates and expanded into three copies two being frozen and a third being digested with proteinase K and the genomic DNA being precipitated, dried and re-suspended in TE for genotyping. ES cell clones were genotyped by PCR across the 5’ (primers APOBEC_1033i2_LR_5F and APOBEC_1033i2_LR_5R) or 3’ (APOBEC-BAC3.5 and APOBEC-BAC3.2) junctions (Suppl. Figure 8A). ES cell clones that had integrated the RP11-1033i2-R38-R39 BAC were tested for completeness by PCR across human specific APOBEC3 A, B, C, D, F, G and H exons (Suppl. Figure 9). To complete the humanization the ES cell clones that had intact BACs were transfected with PBase to remove the selection cassettes from either end of the inserted BAC. ES cells from a single confluent well of a 24 well plate were trypsinized and resuspended, with 1/12 of the volume being passed onto a 24 well plate and the remaining cells being washed with and resuspended in PBS to be electroporated (Biorad) with 10μg of PBase expression plasmid ^40^ and passed to a single well of a 6 well plate. After 3 days culture in M15 the cells were trypsinized and counted and 500,000 cells were plated on a 10cm dish. The following day the media was changed to M15 with 10μM FIAU, and for 10 days subsequently, to select for loss of the *pu tk* gene ^74^ at the 3’ junction. As PBase expression would act on both the selection cassettes derived from pRMCE38 and pRMCE39, their removal was assessed by PCR across the junctions (Suppl. Figure 8B). A selection of the ES cell clones that had passed all humanization steps (Fig 1b) were microinjected into mouse blastocysts to generate male chimearas, which were them bred with C57BL/6 albino females to test for germline transmission. Mice were genotyped using short range or long-range primers, or by loss of allele (LoA) assay which detects if how many wild type alleles remain as an indication of humanized allele zygosity (Suppl. Figure 10).

### Breeding of the Hs-APOBEC3 and mA3 KO (landing pad) lines

Breeding colonies were established at The Wellcome Trust Sanger Institute animal facility. Initial backcrossing to C57BL/6 (5 generations) and breeding was conducted at Charles River Laboratories (Manston, UK) prior to transfer and rederivation by hysterectomy into the Specific Pathogen Free unit at the University of Southampton Biomedical Research Facility, where a further 4 generations of backcrossing to C57BL/6 was performed.

### Mouse embryonic stem cell culture

SNL76/7 feeder cells (ECACC 07032801) were incubated for 3 hours at 37 ᵒC with 10 µg/ml Mitomycin C (Sigma M4287) and washed 3 times with PBS. Treated cells were seeded onto gelatinized plates (0.1% Gelatin, Sigma ES-006) 24 hours prior to plating embryonic stem (ES) cells, with approximately 1.2 x 10^6^ SNL cells added to each T25 plate. Cells were maintained in 5% CO_2_ at 37 ᵒC in media containing DMEM (PAN-Biotech P04-03559), 7% FBS (PAN Biotech P40-37500) and 1X Glutamine-Penicillin-Streptomycin (ThermoFisher 10378016). Mouse ES cells were grown on SNL feeder plates in media containing Advanced DMEM (ThermoFisher 1249105), 16% FBS (Sigma ES-009-B), 1X Glutamine-Penicillin-Streptomycin (ThermoFisher 10378016), 1X Mecaptoethanol (Sigma ES-007-E) and 1000 units/ml Leukaemia Inhibitory Factor (LIF, Millipore ESG1107). Media was replaced every 24 hours. Cells were maintained in 5% CO_2_ at 37 ᵒC and passaged upon reaching 80-85% confluence, approximately every 2-4 days. Media was replaced 3 hours prior to passaging the cells. For harvesting, feeder cells were removed by differential trypsinization and ES cells were collected for RNA or DNA extraction by further trypsinisation and centrifugation at 400 x g. Cell pellets were resuspended in ice-cold PBS and re-centrifuged at 400 x g, followed by removal of PBS flash freezing in liquid nitrogen and storage at −80C.

### PMBC isolation

Whole blood was collected into EDTA tubes from homozygous H-APOBEC3 mice under terminal anaesthesia. PBMCs were isolated from the freshly obtained blood using density gradient separation (Histopaque 1083, Sigma 1083-1), following the manufacturer’s instructions. Typically, 500 µl – 1 ml of blood was collected from each mouse, with the blood from 3 mice pooled for each experiment. Briefly, approximately 3 ml of whole blood was layered onto 3 ml of Histopaque 1083 in a 15 ml centrifuge tube. Each tube was centrifuged at 400 x g for 30 minutes at room temperature. Following centrifugation, the opaque interface containing the PBMCs was transferred to a clean 15 ml centrifuge tube. Cells were resuspended in PBS and centrifuged at 250 x g for 10 minutes. PBS washes were performed 3 times to remove any remaining Histopaque 1083. Following the final wash, cells were resuspended in RPMI 1640 medium (ThermoFisher 11875093) with 10% FBS (PAN Biotech P40-37500). Cells were maintained in 5% CO_2_ at 37 ᵒC.

### Interferon and PMA treatment

Following isolation, PBMCs were incubated in 5% CO_2_ for 24 hours at 37 ᵒC. Media was removed and replaced with one of the following: media alone; media containing 100 ng/ml phorbol ester, phorbol-12-myristate-13-acetate (PMA, Sigma P1585); media containing 1000 units/ml Interferon α from human (Sigma I4276); media containing 1000 units/ml Interferon α from mouse (Sigma I8782). Cells were incubated for 6 hours at 37 ᵒC, following which the media was removed and the cells washed with PBS. All remaining PBS was aspirated off, the cells flash frozen on dry ice and placed at −80 ᵒC for longer term storage.

### RNA isolation from mouse ES cells

RNA was isolated from the mouse ES cells using the Monarch Total RNA Miniprep Kit (NEB T2010), following the manufacturer’s instructions. Briefly, cell pellets were resuspended in 600 µl lysis buffer and loaded onto gDNA removal columns. Columns were centrifuged at 16 000 x g for 1 minute to remove the majority of the gDNA. Following the addition of ethanol, samples were loaded onto RNA purification columns and centrifuged at 16 000 x g for 1 minute. On column treatment with DNAse I was carried out to remove any remaining gDNA and the remaining RNA washed and eluted from the columns with 100 µl nuclease free water. RNA quantification was carried out using a NanoDrop UV-Vis Spectrophotometer (ThermoFisher).

### RNA isolation from PBMCs and tissues

RNA isolation from the PBMCs and mouse tissues (ovary, spleen, lung) was carried out using the Monarch Total RNA Miniprep Kit, following a similar protocol to the mouse ES cells. Prior to the addition of lysis buffer, samples were incubated for 30 minutes at 55 ᵒC with 1X DNA/RNA Protection Reagent, Proteinase K Reaction Buffer, and Proteinase K, at a ratio of 20:2:1. Tissue samples were mechanically homogenized using a handheld grinder (TissueRuptor, Qiagen). Samples were vortexed and centrifuged for 2 minutes at 16 000 x g. The supernatant was transferred to a clean microfuge tube and the lysis buffer added. Following gDNA removal, RNA was washed and eluted using 100 µl nuclease free water and quantification carried out with a NanoDrop UV-Vis Spectrophotometer (ThermoFisher).

### Primer Design

Human *A3A-H* forward and reverse primer sequences were obtained from Refsland *et al.* (2010). *Tbp*, *Cbx6*, *Cbx7* and *Ccne1* primer design was carried out using the IDT Primer Quest Tool (https://eu.idtdna.com/primerquest/home/index). Primer sequences were: *Tbp* forward CTACCGTGAATCTTGGCTGTAA; *Tbp* reverse GTTGTCCGTGGCTCTCTTATT; *Cbx6* forward AAGGGAACGTGAGCTGTATG; *Cbx6* reverse GCTCGGCTTGACAGAGAAA; *Cbx7* forward CTCTCCACCCACATACACTAAC; *Cbx7* reverse TTCAACCCAACCTCTACATTCA; *Ccne1* forward CTGGATGTTGGCTGCTTAGA; *Ccne1* reverse TCTATGTCGCACCACTGATAAC. Primer pair specificity was verified using Primer-BLAST software (https://www.ncbi.nlm.nih.gov/tools/primer-blast). All primers were synthesized by Integrated DNA Technologies (https://eu.idtdna.com).

### qRT-PCR

cDNA synthesis was carried out using the LunaScript RT SuperMix Kit (NEB E3010L), following the manufacturer’s instructions. Up to 1 µg of RNA was used in each 20 µl reaction. Each reaction was placed at 25 ᵒC for 2 minutes, 55 ᵒC for 10 minutes and 95 ᵒC for 1 minute. Quantitative PCR was used to determine gene expression at the RNA level for the different genes of interest. qPCR was carried out using the QuantStudio 3 Real-Time PCR System (ThermoFisher A28567). 10 µl reactions were prepared including 5 µl 2X PowerUp SYBR Green Master Mix (Applied Biosystems A25918), 0.3 µl forward and reverse primers (300 nM each), 2.4 µl H_2_O and 2 µl cDNA (at 5 ng/µl). Reactions were loaded onto 96-well plates (Applied Biosystems N8010560) and sealed with adhesive covers (Applied Biosystems 4311971). No template controls (H_2_O in place of cDNA) were included for each set of primers on each qPCR plate and technical replicates were included for each reaction. Reactions were placed at 50 ᵒC for 2 minutes, 95 ᵒC for 10 minutes and then 40 cycles of 95 ᵒC for 15 seconds and 60 ᵒC for 1 minute. Following the 40 cycles, the temperature was ramped from 60 ᵒC to 95 ᵒC at a rate of 0.15 ᵒC/second for melt curve analysis. In the analysis of hA3, *Mdm2* and *Ckdn1a* (p21) in Figure 2, gene expression was measured using the following Taqman assays (ThermoFisher): *APOBEC3A* (Hs00377444); *APOBEC3B* (Hs00358981); *APOBEC3C* (Hs00828074); *APOBEC3D* (Hs00537163); *APOBEC3F* (Hs01665324); *APOBEC3G* (Hs00222415); *APOBEC3H* (Hs00419665); *Mdm2* (Mm00519571); *Cdkn1a* (Mm01268809). Biological replicates were performed for each sample, with a minimum of n=3 biological repeats.

### ChIP-qPCR methods

10^7^ mouse ES cells (clone 1A1_11) were used per ChIP experiment. After 24hrs treatment, cells were fixed with 1% formaldehyde for 10 min at 37°C. Crosslinking was quenched by adding glycine to a final concentration of 0.1 M. Cells were lysed in LB1 buffer (50 mM HEPES-KOH (pH 7.5), 140 mM NaCl, 1 mM EDTA, 10% glycerol, 0.5% NP-40 and 0.25% Triton X-100) followed by mixing on rotator for 10 min at 4 °C. Then, nuclei pellets were collected and resuspended in LB2 buffer (10 mM Tris-HCL (pH 8.0), 200 mM NaCl, 1 mM EDTA and 0.5 mM EGTA) and mixing on rotator at 4 °C for 10 min. Then, pellets were collected and resuspended in LB3 buffer (10 mM Tris-HCl (pH 8), 100 mM NaCl, 1 mM EDTA, 0.5 mM EGTA, 0.1% Na-deoxycholate and 0.5% N-lauroylsarcosine). Chromatin was sonicated (Diagenode) to get DNA fragments of 100–1,000 bp size. Sonicated lysates were incubated overnight at 4 °C with 100 μl of Dynabeads® Protein G (Invitrogen) and 5 μg of antibody (p53 antibody, sc-126, Santacruz). The next day the beads were collected and washed 6 times with 1 ml ice-cold RIPA buffer and twice with TBS (pH 7.4). Both ChIP and input samples were reverse crosslinked by adding 200 μl of elution buffer (1% SDS, 0.1 M NaHCO3) and incubating overnight at 65°C. Following reverse crosslinking, DNA was purified using the phenol-chloroform-isoamyl alcohol extraction method. The purified DNA was then used for qPCR. ChIP qPCR primers are listed in Suppl. Figure 11.

### Statistical Analysis

Standard curves for all primer pairs were generated using plasmids containing the relevant regions of cDNA, using 10-fold dilution series ranging from 10^1^ to 10^8^ copies. *A3, Cbx6, Cbx7 and Ccne1* expression data were normalized to the expression data for *Tbp* for each sample, shown as copy number relative to *Tbp*. All data points are shown as mean ± SEM of the biological repeats.

### Aging Study

Hs-APOBEC3 littermates of each genotype (WT, HET, HOM) and *Apobec3* KO mice were maintained for up to 730 days to determine whether there was any difference in survival probability between the groups (WT n=21; HET n=26; HOM n=25; KO n=10). Weight and length (nose to base of tail) measurements were taken for each mouse that survived to 365 days of age.

### F-MLV infection of splenocytes

Spleens were harvested from adult littermates that were either WT, heterozygous or homozygous for the Hs-APOBEC3 locus and were transported in HEPES-buffered Iscove’s Modified Dulbecco’s Medium (PAN Biotech P04-20150). Spleens were strained through a 100 micron cell strainer (Fisherbrand Cat. No. 22363549)) into 5 mL of RPMI (Gibco 21875-034). The resulting cell suspension was centrifuged at 370g for 7 minutes at 10°C and the pellet resuspended in 2 mL RBC lysis buffer (eBioscienceTM 00-4300) for 2 minutes at room temperature before a further 13 mL of RPMI was added. Cells were then centrifuged again for 7 minutes at 10°C, and the pellet resuspended in RPMI to a concentration of 3 x 10^6^ cells per mL. LPS (Enzo ALX-581-008) was added to a concentration of 10 ug/ml and cells were incubated at 37°C. Two days later, 3 x 10^6^ splenocytes were pelleted and resuspended in 3 mL of either RPMI (mock control) or virus-containing, filtered culture supernatant from BHF cells (from Dr. George Kassiotis) and 3 ug/mL Polybrene (SIGMA H9268-5G). BHF cells are a *Mus dunni* tail fibroblast (MDTF) cell line that is chronically infected with Friend murine leukaemia virus (F-MLV). The splenocytes were incubated for 3 days at 37°C before pelleting and resuspending in 3 mL of fresh RPMI.

### MDTF infections

MDTF cells were seeded in 12-well plates at a density of 5 x 10^4^ cells per well. The following day, cells were pre-treated with 10 ug/mL polybrene for 1 hour before the media was removed and replaced with 1 mL of the infected splenocyte suspension or uninfected splenocytes as a control. After 24 h, the MDTF cells were washed twice with pre-warmed DMEM (Gibco 41966-029) and incubated with fresh DMEM. Samples were then analysed in multiple ways: After 2 days and 5 days, 0.15 mL of culture supernatant was harvested and viral RNA was extracted using QIAamp Viral RNA mini Kit (Qiagen). The number of genomic RNA copies was quantified using TaqMan™ Fast Virus 1-Step Master Mix with a primer set amplifying the env gene of Fr-MLV (For: CTGCGCCAGAGACTGCGACGA Rev:GACCCGGGGCAGACATAAAAT) (Thorborn et al 2014). At 5 days, cells were fixed with 4% paraformaldehyde and immunostained with anti-F-MLV glycogag antibody (from Dr. George Kassiotis, 1:500 dilution) followed by Goat anti-mouse IgG (H+L) HRP conjugate at 1:1000 (Biorad 172-1011). The number of infected cells were revealed using True Blue™ peroxidase substrate (KPL 50-78-02) supplemented with 0.03% H2O2 (1:1000 of 30% solution SIGMA H1009-100ml). In parallel, 5-day cell cultures were harvested and total genomic DNA was isolated using the DNeasy blood and tissue kit (QIAGEN 69506). This DNA was sequenced and analysed for mutations.

### F-MLV *env* sequencing

Amplicons of 2.186 kb corresponding to the env and U3 region of F-MLV were generated using primers that were modified from Salas-Briceno and Ross ^62^. The initial 5 cycles of PCR were performed using Q5 polymerase (NEB M0491), 200uM dNTPs (Invitrogen 10297-018) and 500nM each of the forward primer (5’-TTTCTGTTGGTGCTGATATTGCNNNYRNNNYRNNNYRNNNccaatggggatcgggagac ag-3’) and reverse primer (5’-ACTTGCCTGTCGCTCTATCTTCNNNYRNNNYRNNNYRNNNgggtcttaagaaactgctga gg-3’, where N indicates random bases that make up unique molecular identifiers (UMIs) for each PCR reaction ^75^ and lower-case letters represent sequence that anneals to the viral sequence. An initial denaturation step of 98°C for 2 minutes was used, followed by 5 cycles of denaturation at 98°C for 10 seconds, annealing at 66°C for 30 seconds and extension at 72°C for 2 minutes. The products were purified using QIAquick PCR purification kit (QIAGEN 28106) before a second PCR reaction was performed with primer pair 5’-TTTCTGTTGGTGCTGATATTGC-3’ and 5’-ACTTGCCTGTCGCTCTATCTTC-3’, using an initial denaturation step at 98°C for 2 minutes, followed by 35 cycles of denaturation at 98°C for 10 seconds, annealing at 60°C for 30 seconds and extension at 72°C for 2 minutes. A final extension was performed at 72°C for 2 minutes. The products were gel purified using QIAquick gel extraction kit (QIAGEN 28706) and sent for Oxford Nanopore sequencing (Full Circle).

### Analysis of FMLV *env* sequencing data

Fastq reads were combined across runs per sample and filtered by length select reads long enough to contain the full reference sequence and both UMIs and some or all of the adapter sequences, to exclude chimeric reads (min=2222, max=2266) using seqkit (v2.1.0). Reads were mapped to the reference sequence for the FMLV amplicon with minimap2 (v2-2.24) and sorted and indexed with samtools (v.1.16.1). 3’ and 5’ UMI sequences were extracted as 18nt from the start and end of mapping. These 3’ and 5’ sequences were concatenated to provide a combined 36nt UMI. Only reads with complete UMIs were included, and any duplicate UMIs reads were removed keeping one read from each UMI (samtools v1.16.1 and custom code). Variant sites were called with freebayes (v1.3.6), using the pooled-continuous option, and a minimum alternate fraction of 0.01. Only variant sites of type “snp” were kept, and haplotype alleles were normalised to individual sites using bcftools norm (1.16). G site locations and their context (1nt following) were extracted from the reference sequence generating bed files (seqkit v2.1.0). Pileups were generated for these sites per read using samtools (v1.16.1), and counts of G sites mutated to A per read were generated per sample. Counts of variant sites, variant allele frequency, and G>A counts per read were summarised, analysed and plotted in Rstudio using R/4.3.2 using the following packages, tidyverse 2.0.0, ggplot2 3.4.4, ggpubr 0.6.6.

## Data Availability

RNA-seq data from H1 ESCs was from GEO accession GSE75297. RNA-seq data from PBMCs from healthy human controls was from GEO accession GSE115262. RNA-seq data for human lung, spleen and ovary was from the Genotype Tissue Expression (GTEx) portal (https://www.gtexportal.org/home/). F-MLV sequencing data will be deposited at GEO upon publication.

## Author Contributions

T.R.F., S.A., J.S., H.M.P, A.B. conceived the study; T.R.F, S.A., J.S. obtained funding; T.R.F., S.A., N.K.K., K.B., M.Y., S.G.C., H.M.P., A.B. designed experiments; H.M.P., R.C., N.K.K., M.P., M.Y. performed experiments; A.S., E.H., N.Z. provided technical assistance; N.K.K., I.G.R., M.P., J.G., G.J.T. performed data analysis, N.K.K., H.M.P, T.R.F. wrote the manuscript., S.A., K.B., M.P., S.G.C., G.J.T., H.M.P., J.S. reviewed the manuscript.

## Acknowledgments

We thank Drs. George Kassiotis and Jonathan Stoye for reagents and advice on F-MLV infection of splenocytes. This work was supported by grants to T.R.F from Cancer Research UK (A25825), BBSRC (BB/V010271/2), The Oddfellows Society H. A. Andrews Memorial Fund and the University of Southampton Centre for Cancer Immunology. We are grateful for expert technical assistance and advice from the Biomedical Research Facility, Research Histology Facility and Biomedical Imaging Unit at the University of Southampton and the animal unit at The Wellcome Trust Sanger Institute. K.N.B is funded by the Francis Crick Institute, which receives its core funding from Cancer Research UK (CC2056), the UK Medical Research Council (CC2056) and the Wellcome Trust (CC2056). For the purpose of Open Access, the author has applied a CC BY public copyright licence to any Author Accepted Manuscript version arising from this submission.

