## Supplementary material for "Human-like *APOBEC3* gene expression and anti-viral responses following replacement of mouse *Apobec3* with the 7-gene human *APOBEC3* locus": Suppl. Figure 1

### Supplementary Figure 1

**A**

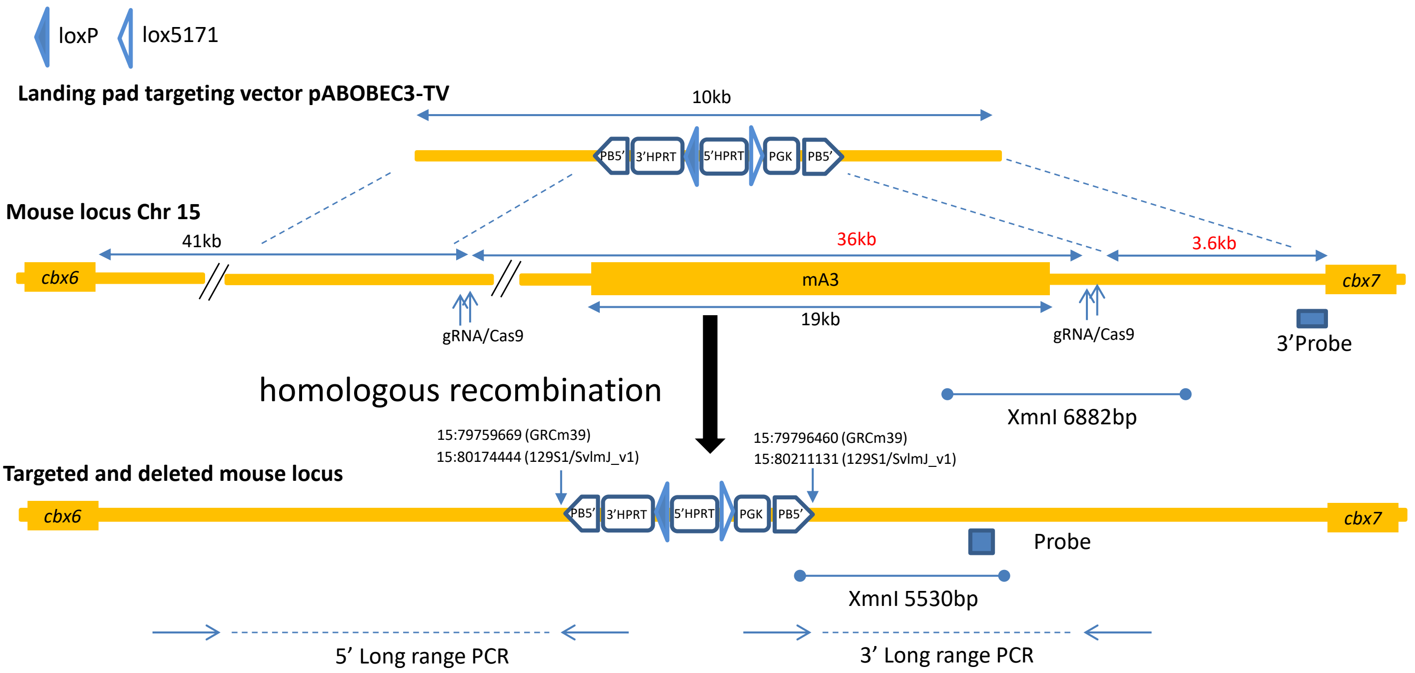


**B**

i) Southern blot using 3' probe against XmnI digest

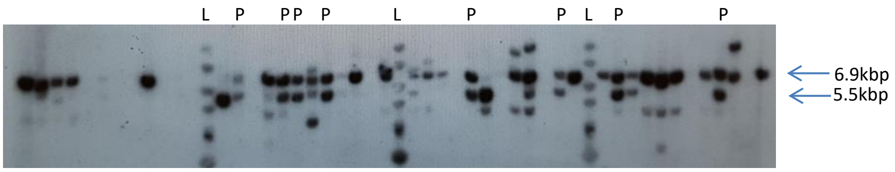


ii) PCR to detect landing pad integration at 3' end

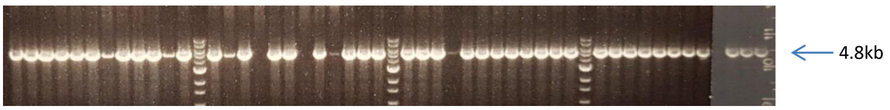


iii) PCR to detect landing pad integration at 5' end

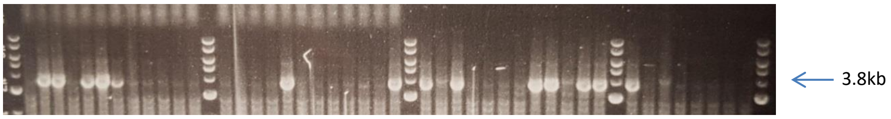


iv) PCR primers for genotyping landing pad

| Assay | Primer name | Primer sequence | Product size |
|---|---|---|---|
| MUT LR PCR 5' | APOBEC PCR EL8<br>APOBEC PCR EL1 | agataaacccaggtttgactcccaaa<br>atgttctagttctgtggccatctgc | 3833 bp |
| MUT LR PCR 3' | APOBEC 3 LR F<br>APOBEC 3 LR R | taggtatgcaaaataaatcaaggtcataac<br>tactgatattggtaaaagactgtatagcat | 4815 bp |

**Supplementary Figure 1: Replacement of *Apobec3* with a landing pad to allow insertion of the Hs-APOBEC3 locus. (A)** Schematic illustrating the steps used for humanizing the Apobec3 locus. A "landing-pad" cassette was targeted by homologous recombination in mouse ES cells, deleting mouse APOBEC3 between 15: 79759669 to 79796460 (GRCm39) which is equivalent to 15:80174444 to 80211131 for 129S1/SvlmJ_v1. Human BAC clone RP11-1033i2 modified and trimmed by flanking cassettes to contain sequence between 22:38921158 to 39130175 (GRCh38), was co-transfected with a Cre expression plasmid insertion into the landing pad loxP and lox-5171 sites by RMCE was selected using Puromycin followed by G418. Finally the flanking cassettes were excised from the locus by transfection with a PBase expression plasmid and selection with FIAU. **(B)** Summary of relation between ES cell clones and mouse colonies for the sequential steps in humanizing the mouse APOBEC gene.
