## Supplementary material for "Human-like *APOBEC3* gene expression and anti-viral responses following replacement of mouse *Apobec3* with the 7-gene human *APOBEC3* locus": Suppl. Figure 2

### Supplementary Figure 2

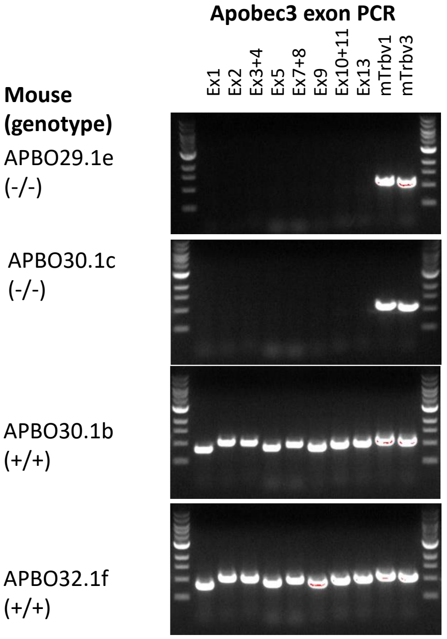


| Primer names | Forward | Reverse | Product |
|---|---|---|---|
| mApobec3-Ex1 | cacaggatggtctaggcaggttg | cctcttctcccagtatcccctga | 847 |
| mApobec3-Ex2 | cctgcttggaaaggactccaga | cgcaggcctttggtctttattg | 1083 |
| mApobec3-Ex3+4 | catccacgctgaaatctgctttt | gaccatctagaggccaagggaga | 1057 |
| mApobec3-Ex5 | agggctaagaggcatctttgtgg | tgtccatggccaacctatgaact | 917 |
| mApobec3-Ex7+8 | acttgccaaagcaggctgaaac | ctgggaaagtccctggtttgtg | 1001 |
| mApobec3-Ex9 | ggggagtagccctggctagaga | aggtgcccagagaaaagggttc | 1079 |
| mApobec3-Ex10+11 | agatagtgcccctggtctccaag | aggaagatgggcaacaggaatgt | 1015 |
| mApobec3-Ex13 | actggactggggaacttcaaagc | agtctcccaagtggcttttggtc | 1051 |
| mTrbv1 | gtggctacaggtcagatggaaaatg | gaggtcaatgtgggtgtgggtaata | 1153 |
| mTrbv3 | aaacctcagaccacaataccacctg | caacagggcagtgaccctacttcta | 1120 |

**Supplementary Figure 2: Verification of Apobec3 deletion in landing-pad mouse line APBO.** To establish that the mouse Apobec3 gene had been properly deleted by insertion go the landing pad cassette, PCR was conducted on genomic DNA PCR (tail) from two homozygous APBO mice (APBO29.1e and APBO29.1c) and two wild type controls (APBO30.1b and APBO32.1f). Homozygous mice lacked all Apobec3 exons tested whereas the wild type mice retained all exons. The table shows the sequences of the mouse Apobec3 exon flanking primers that were used for the PCR together with two control primers (mTrbv1 and mTrbv3) from a different locus. The ES cell clone from, which APBO mice were derived, were later used for integration of the modified RP11-1033i2 BAC clone.
