## Supplementary material for "Human-like *APOBEC3* gene expression and anti-viral responses following replacement of mouse *Apobec3* with the 7-gene human *APOBEC3* locus": Suppl. Figure 3

### Supplementary Figure 3

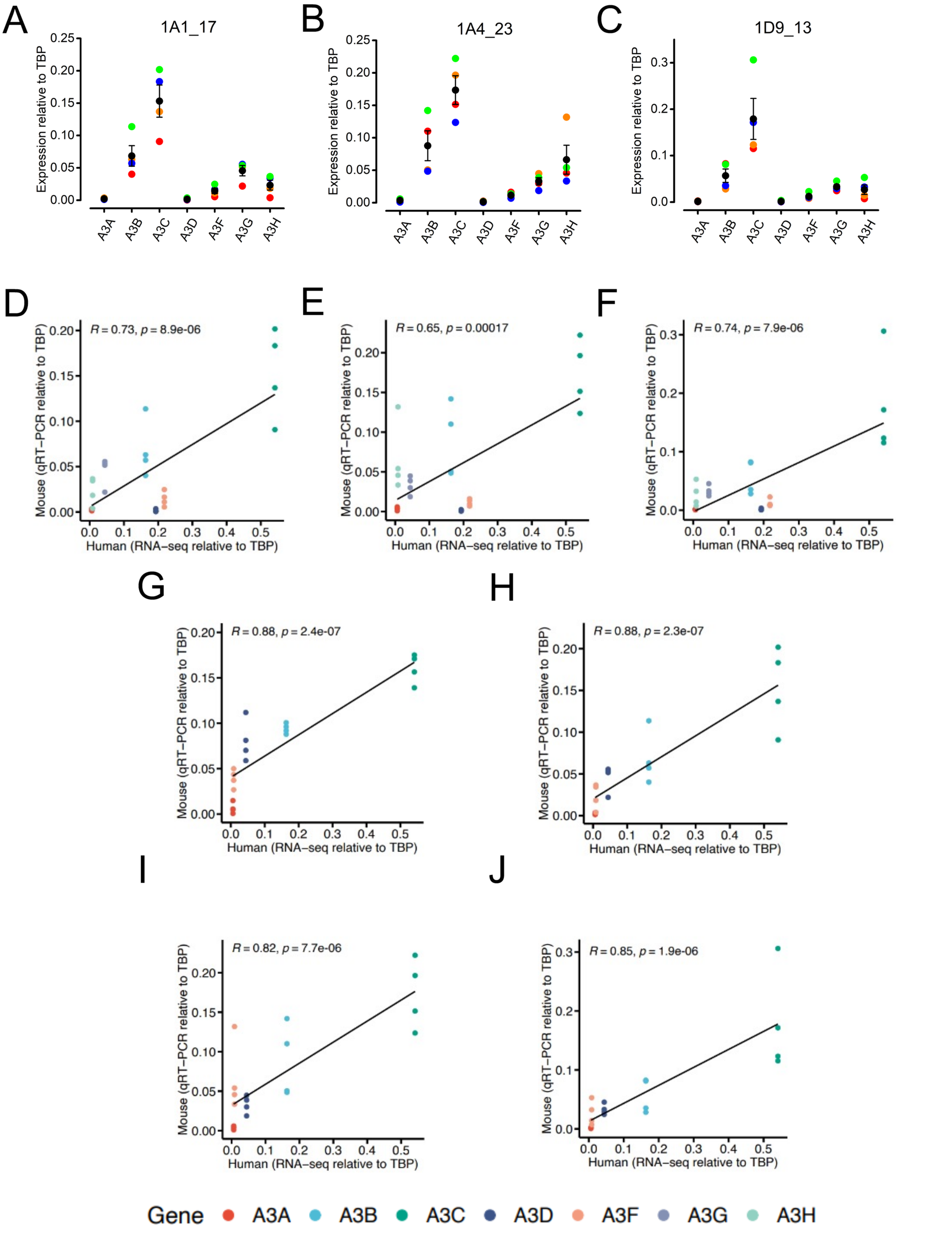


**Supplementary Figure 3: Correspondence between hA3 gene expression in human and Hs-APOBEC3 across three additional mouse ES cell clones. (A-C)** hA3 gene expression levels in ES cell clones 1A1_17 **(A)**, 1A4_23 **(B)** and 1D9_13 **(C)**. Expression levels are normalized to the reference gene *Tbp* (n=4; red, repeat 1; orange, repeat 2; blue, repeat 3; green, repeat four; black, mean). **(D-F)** hA3 gene expression in mouse ES cell clones 1A1_17 **(D)**, 1A4_23 **(E)** and 1D9_13 **(F)** compared with RNA-seq data from human ES cell clone H1. **(G-J)** comparisons between hA3 gene expression in mouse ES cell clones 1A1_11 **(G)**, 1A1_17 **(H)**, 1A4_23 **(I)** and 1D9_13 (J) and human ES cell clone H1 with A3D and A3F omitted. Accompanies Figure 2.
