## Supplementary material for "Human-like *APOBEC3* gene expression and anti-viral responses following replacement of mouse *Apobec3* with the 7-gene human *APOBEC3* locus": Suppl. Figure 4

### Supplementary Figure 4

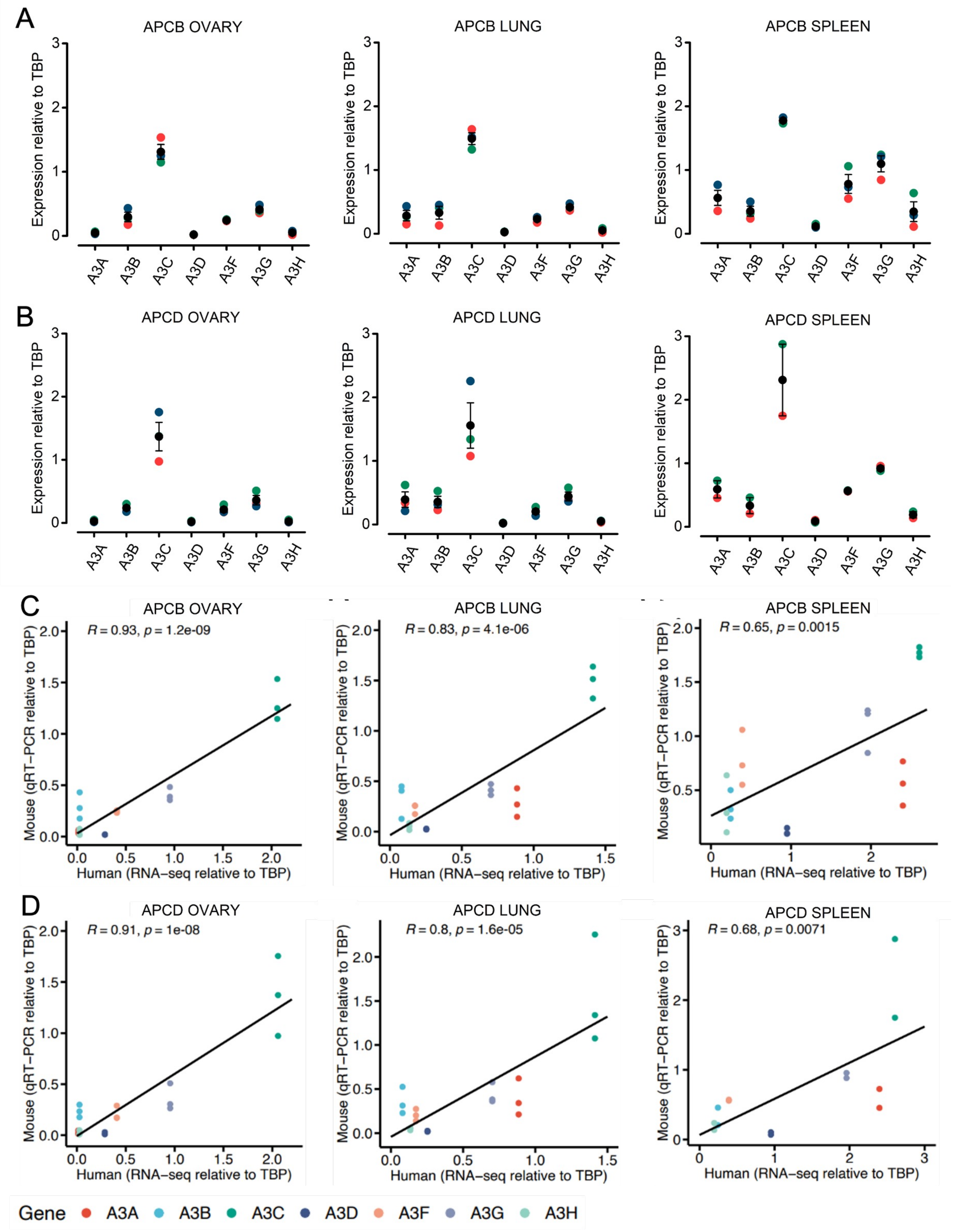


**Supplementary Figure 4. Tissues from Hs-APOBEC lines APCB and APCD express hA3 genes at comparable levels to equivalent human tissues. (A, B)** hA3 gene expression levels in the ovary (left panels), lung (middle panels) and spleen (right panels) of adult homozygous APCB **(A)** and APCD **(B)** mice. RT-qPCR data are shown relative to *Tbp* (n=3; mouse 1, red; mouse 2, orange; mouse 3, green; mean, black). Error bars show SEM. **(C, D)** hA3 gene expression in the tissues from APCB **(C)** and APCD **(D)** mice compared with the corresponding human data from the Genotype-Tissue Expression (GTEx) Portal for the ovary (left panels), lung (middle panels) and spleen (right panels). Pearson correlation coefficients are indicated for each tissue.
