## Supplementary material for "Human-like *APOBEC3* gene expression and anti-viral responses following replacement of mouse *Apobec3* with the 7-gene human *APOBEC3* locus": Suppl. Figure 5

### Supplementary Figure 5

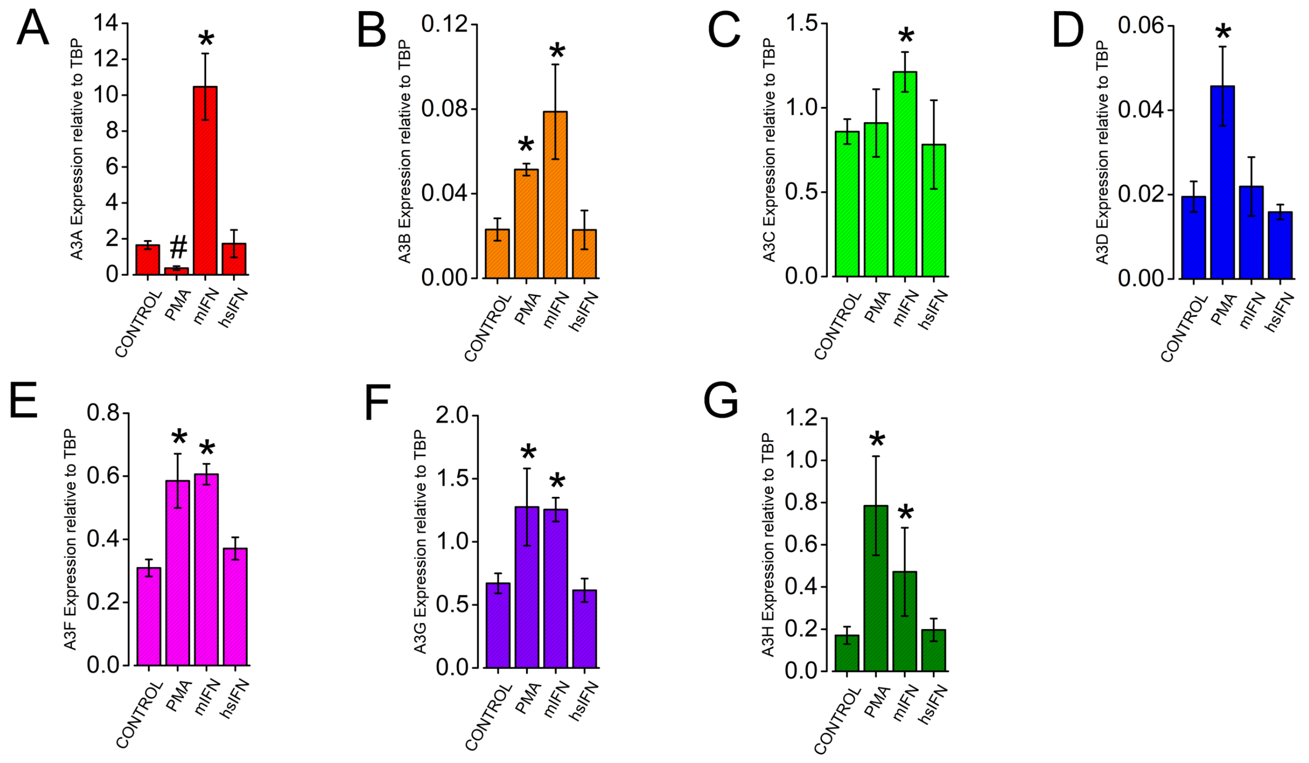


**Supplementary Figure 6: Individual plots representing the hA3 gene expression data displayed in Figure 4C.** hA3 gene expression levels in the humanized mouse PBMCs following 6-hour exposure to control media (n=6), PMA (n=4), mouse IFNα (n=3) or human IFNα (n=3) shown for all 7 hA3 genes. Error bars show SEM. Accompanies Figure 4.
