## Supplementary material for "Human-like *APOBEC3* gene expression and anti-viral responses following replacement of mouse *Apobec3* with the 7-gene human *APOBEC3* locus": Suppl. Figure 6

### Supplementary Figure 6

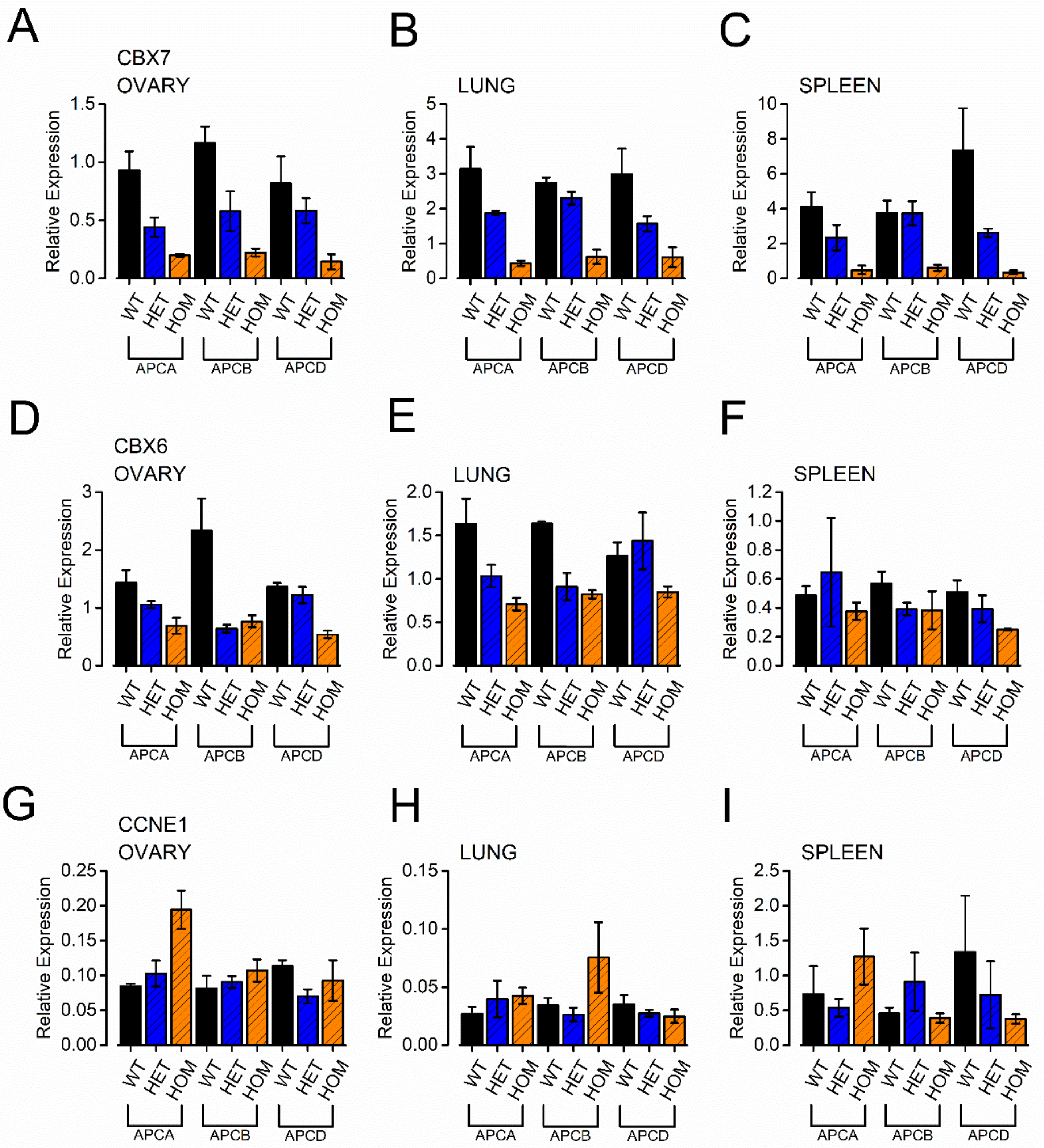


**Supplementary Figure 6:** qRT-PCR measurements of *Cbx7* (A-C), *Cbx6* (D-F) and *Ccne1* (G-I) expression in ovary (A, D, G), lung (B, E, H) and spleen (C, F, I) from the three Hs-APOBEC3 mouse lines (APCA, APCB, APCD). All gene expression measurements are normalised to *Tbp*. Error bars = SEM. Accompanies Figure 5.
