## Supplementary material for "Human-like *APOBEC3* gene expression and anti-viral responses following replacement of mouse *Apobec3* with the 7-gene human *APOBEC3* locus": Suppl. Figure 7

### Supplementary Figure 7

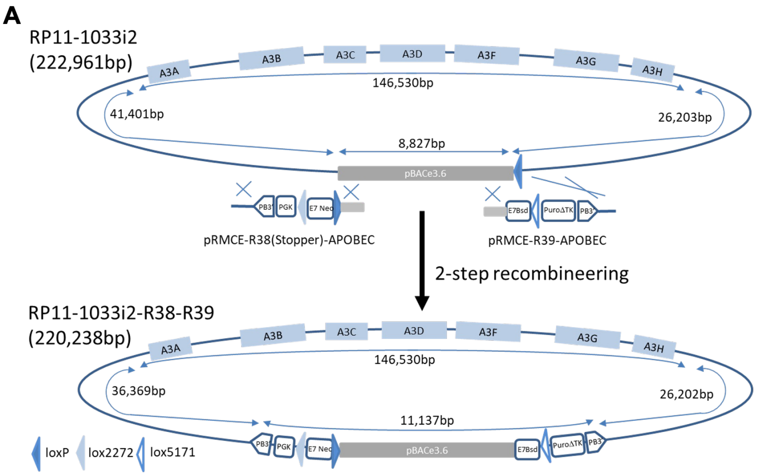


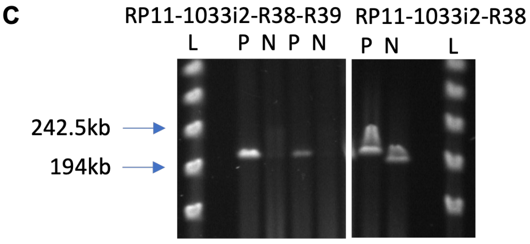


| BAC clone/Digest | PmeI | NotI |
|---|---|---|
| RP11-1033i2-R38-R39 | 220,238bp | Uncut |
| RP11-1033i2 | 222,961bp | 214,227+8,734bp |

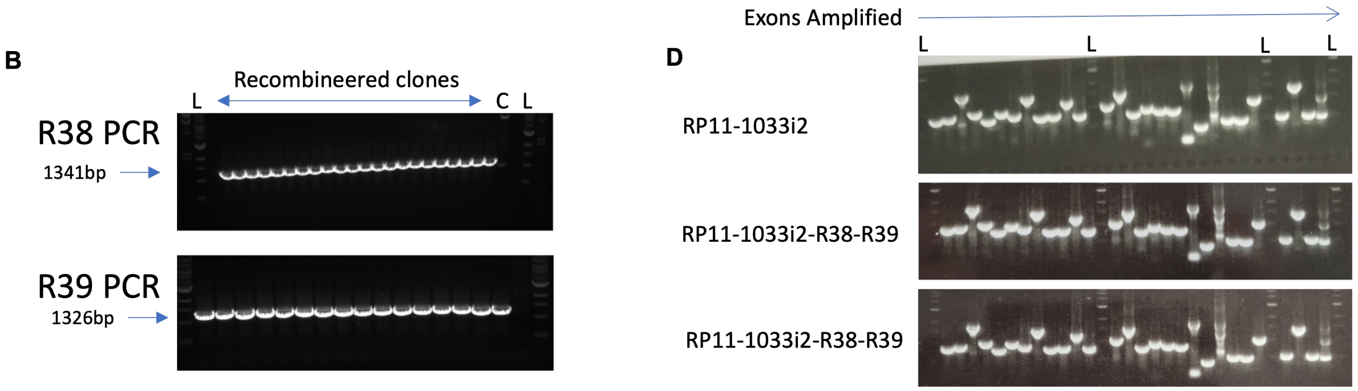


**Supplementary Figure 7: Preparation of human BAC RP11-1033i2 for transgenesis. (A)** Recombineering of BAC RP11-1033i2. Human BAC RP11-1033i2 was modified by recombineering in *E.coli* using vectors pRMCE38 and pRMCE39 (Li et al, 2014). The pRMCE38 had previously been modified as described in supplementary materials methods. **(B)** PCR verification of recombineered BAC ends. The correct integration of recombineering cassettes was verified by PCR extending from BAC internal sequences to the cassettes sequences. Controls are marked as C where the template was wild type RP11-1033i2 BAC, which gave no specific amplifying products. Lanes containing 1kb ladder are marked L. **(C)** Pulse Field Gel Electrophoresis (PFGE) of BAC clones. To ensure that modified BACs were of the correct size, and did not contain gross rearrangements, they were digests with PmeI (P) or NotI (N) and analysed by PFGE. The two RP11-1033i2-R38-R39 clones shown here appeared to be the correct (expectations summarized in the table), linearizing with PmeI and remaining uncut with NotI. Lanes containing Lambda PFG Ladder (NEB) are marked L. **(D)** Integrity analysis of BAC clones by PCR. Recombineered BAC clones were further demonstrated as intact by PCR across APOBEC3 exons using the same primers listed in Suppl. Fig 5. The WT RP11-1033i2 clone and two RP11-1033i2-R38-R39 clones contained all the expected exon sequences. Lanes containing 1kb ladder are marked L.
