## Supplementary material for "Human-like *APOBEC3* gene expression and anti-viral responses following replacement of mouse *Apobec3* with the 7-gene human *APOBEC3* locus": Suppl. Figure 8

### Supplementary Figure 8

**A**

5’ Long
APOBEC_1033i2_LR_5’

3’ Long
APOBEC_1033i2_LR_3’

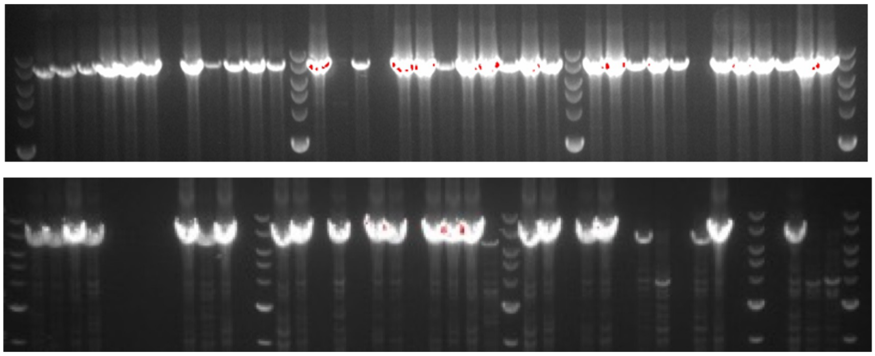

7689 bp

6622 bp

**B**

5’ Long
APOBEC_1033i2_LR_5’

3’ Long
APOBEC_1033i2_LR_3’

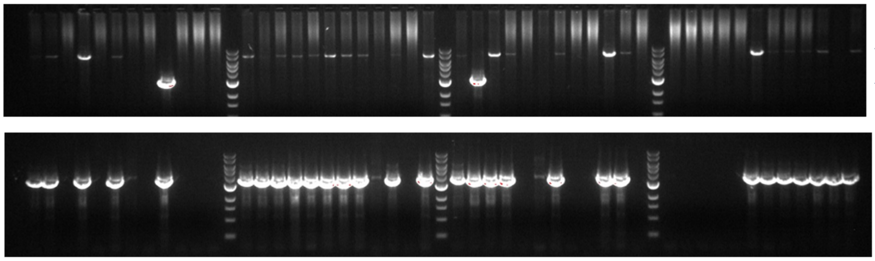

7689 bp
(Undeleted)

3087 bp
(Deleted)

3562 bp
(Deleted)

**C**

| PCR | Primer name | Primer sequence | Pre PB Product | Post PB Product |
|---|---|---|---|---|
| APOBEC_1033i2_LR_5’ | APOBEC_1033i2_LR_5F<br>APOBEC_1033i2_LR_5R | TAGTTCACTACCACCTATAACTCCAACTT<br>GTAAGTACTTTAAAGCATGTCTACAAATCC | 7689bp | 3087bp |
| APOBEC_1033i2_LR_3’ | APOBEC-BAC3.5<br>APOBEC-BAC3.2 | TGTTTAACTCTTCTCCCCACCATCA<br>CCACTGCTCTACTGAGCTCAACTCC | 6622bp | 3562bp |

**Supplementary Figure 8: Integration of human BAC RP11-1033i2. (A)** PCR for integration of modified RP11-1033i2 BAC. Integration of the modified RP11-1033i2 BAC clone carrying the human APOBEC3 locus into the landing pad integration site of ES cell clone Apobec#A03-2D was assessed by PCR across the 5’ and 3’ junctions of transfected and selected clones. **(B)** PCR for detecting deletion of selection cassettes from BAC ends. Following PiggyBAC transfection and selection for loss of the 3’ selection cassette, ES cell clones were assessed for loss of the 5’ and 3’ selection cassettes by PCR. **(C)** PCR primers used for detecting BAC integration and selection cassette removal. The sequences of long range PCR primers used for the analysis are shown.
