## Supplementary material for "Human-like *APOBEC3* gene expression and anti-viral responses following replacement of mouse *Apobec3* with the 7-gene human *APOBEC3* locus": Suppl. Figure 9

### Supplementary Figure 9

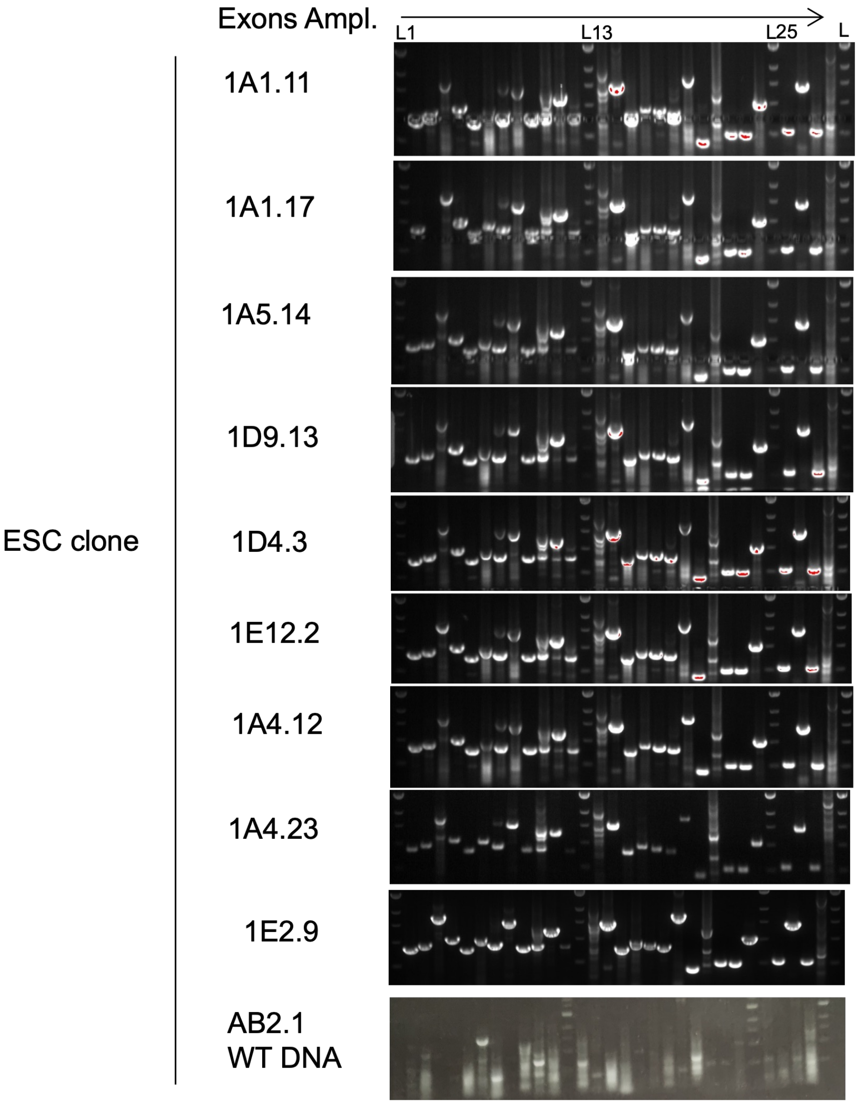

| Lane | Primer Name | Sequence | Product |
|---|---|---|---|
| 1 | A3A Exon1 F | aagtgtggccagctctacat | 871 |
|  | A3A Exon1 R | cgagtagacctgggcatgaa |  |
| 2 | A3A Exon2 F | gctcggtgtggtaggagtta | 932 |
|  | A3A Exon2 R | tctcctgcctcagtctctct |  |
| 3 | A3A Exon3-5 F | tgtgtatagtggagctgggg | 1761 |
|  | A3A Exon3-5 R | gggttcacgctactctcctg |  |
| 4 | A3B Exon1 F | cccactggagtctcaaaacc | 1041 |
|  | A3B Exon1 R | ggacatctgctcaccatgga |  |
| 5 | A3B Exon2 F | cccgctcagcacttagaaga | 802 |
|  | A3B Exon2 R | cgcctttgttctgcttctgt |  |
| 6 | A3B Exon3-4 F | ggacaagaggaagcaggagt | 1009 |
|  | A3B Exon3-4 R | agcaggtgagaggagagaga |  |
| 7 | A3B Exon5 F | aatgctgggtgtggtaggag | 900 |
|  | A3B Exon5 R | tggccttgtgagtcaggtag |  |
| 8 | A3B Exon6-8 F | ctgtgtatattggagctgggg | 1506 |
|  | A3B Exon6-8 R | ctttgctggtgtctgtgagc |  |
| 9 | A3C Exon1 F | agcagcctgtctttatcgga | 814 |
|  | A3C Exon1 R | ggtcgtgtgcatcctactct |  |
| 10 | A3C Exon2 F | gagaatcccttgaacccgag | 841 |
|  | A3C Exon2 R | aaagaattggctgggcatgg |  |
| 11 | A3C Exon3-4 F | atcccagcttatcaggaggc | 1244 |
|  | A3C Exon3-4 R | acgcaattctcccacctttg |  |
| 12 | A3D Exon1 F | cgcttcccaggttcaagtg | 800 |
|  | A3D Exon1 R | cagagaccattccctcccac |  |
| 13 | A3D Exon2 F | ccacgcagcacttagaagag | 1079 |
|  | A3D Exon2 R | tataggaagtggccaggcac |  |
| 14 | A3D Exon3-4 F | cagaattcacacacgaggcc | 1462 |
|  | A3D Exon3-4 R | gcctctcactcacaggaaga |  |
| 15 | A3D Exon5 F | atgggttgggagatgtgtgg | 808 |
|  | A3D Exon5 R | atcctagaacgttgggaggc |  |
| 16 | A3D Exon6-7 F | ctacactcctcctgctcctg | 943 |
|  | A3D Exon6-7 R | ctggggagatgggaagagtc |  |
| 17 | A3F Exon1 F | acacctggccctttactctc | 872 |
|  | A3F Exon1 R | agagttcaggaatggcggaa |  |
| 18 | A3F Exon2 F | tgagccataatgtccagcca | 835 |
|  | A3F Exon2 R | ctcgggagtcttaggcgac |  |
| 19 | A3F Exon3 F | tgtagtgagccgagattgca | 1739 |
|  | A3F Exon3 R | aagaaggggaaggagcacaa |  |
| 20 | A3F Exon4 F | atcagctccgaggaatccag | 407 |
|  | A3F Exon4 R | gagcagagatcacgccattg |  |
| 21 | A3F Exon5-6 F | aaaattaggcaggcgtggtg | 1229 |
|  | A3F Exon5-6 R | gggtgacagagcgaaattcc |  |
| 22 | A3G Exon1 F | ggaggtcactttagggaggg | 513 |
|  | A3G Exon1 R | ccatcaaagccctgtgaaca |  |
| 23 | A3G Exon2 F | agtctccgccgaaattctca | 509 |
|  | A3G Exon2 R | acggaaacacaaggcctttt |  |
| 24 | A3G Exon3-4 F | atagctgcttgtaggtgcca | 1039 |
|  | A3G Exon3-4 R | tgtgggtgagggagtgtaac |  |
| 25 | A3G Exon5 F | tctagcaagtgagtgggagc | 553 |
|  | A3G Exon5 R | ggtgttagcccctgagacat |  |
| 26 | A3G Exon6-8 F | tttgagaccagcctgaacca | 1449 |
|  | A3G Exon6-8 R | gctttgctggtgtctgtgat |  |
| 27 | A3H Exon1 F | cttgaacttggaaggcggag | 547 |
|  | A3H Exon1 R | acgggaaagaatgggagtgt |  |
| 28 | A3H Exon2-4 F | ctctctcttctcccctccct | 1698 |
|  | A3H Exon2-4 R | gagtcttgctttgttgccca |  |

**Supplementary Figure 9: PCR across APOBEC3 exons for modified RP11-1033i2 BAC transgenic ES cell clones.** ES cell clones that had integrated the modified RP11-1033i2 BAC clone were assessed to determine if the human DNA was intact by PCR across the APOBEC 3 exons using the primers in the table on the right. Note that the four of the primer pairs (A3C Exon2, A3D Exon2, A3F Exon5-6 and A3H Exon2-4) gave a high level of non-specific amplification in all cases, and so were not useful for this analysis). Wild type AB2.1 DNA gave no specific amplification products. Lanes containing 1kb ladder are marked L.
