## Supplementary material for "Human-like *APOBEC3* gene expression and anti-viral responses following replacement of mouse *Apobec3* with the 7-gene human *APOBEC3* locus": Suppl. Figure 10

### Supplementary Figure 10

**A**

| Assay | Primer name | Primer sequence | Product size | T anneal (°C) |
|---|---|---|---|---|
| APOBEC_1033i2_LR_5’ | APOBEC_1033i2_LR_5F<br>APOBEC_1033i2_LR_5R | TAGTTCACTACCACCTATAACTCCAACTT<br>GTAAGTACTTTAAAGCATGTCTACAAATCC | 3087 bp | 55 |
| APOBEC_1033i2_LR_3’ | APOBEC-BAC3.5<br>APOBEC-BAC3.2 | TGTTTAACTCTTCTCCCCACCATCA<br>CCACTGCTCTACTGAGCTCAACTCC | 3562 bp | 60 |
| WT LR PCR | APOBEC WT LRPCR F<br>APOBEC WT LRPCR R | CTGAACATAGGAAATAAACAAATCTTCTGT<br>ATTCTTTCAGTTTCTTCAATCTATTTCCTG | 6432 bp (C57B/6)<br>3298 bp (129) | 60 |
| MUT SR PCR 3’ | APOBEC SRPCR 3.1<br>APOBEC SRPCR 3.2 | CATGACAGTCCTGCGATCCGGAATC<br>GGGTGGGTGTTTCTGGGGTGAATAA | 402 bp | 60 |
| WT SR PCR | APOBEC WT F<br>APOBEC WT LRPCR R | TGGAAGATTTATTACAGATTTCTCTACCAA<br>ATTCTTTCAGTTTCTTCAATCTATTTCCTG | 1182 bp | 60 |

**B**

| LoA Assay | Taqman Code | Forward Primer | Reverse Primer | Probe |
|---|---|---|---|---|
| Apobec3_HP_WT | APNKT7D | TATGTCCTGGAGCCCCT GTT | TGGTGTGTAGCCAGGAAC CTT | CGAATGTGCAGAGCAGA |

**Supplementary Figure 10: Summary primers suitable for genotyping mice. (A)** Long- and short-range PCR primers. **(B)** Apobec3 LoA assay.
