## Supplementary material for "Human-like *APOBEC3* gene expression and anti-viral responses following replacement of mouse *Apobec3* with the 7-gene human *APOBEC3* locus": Suppl. Figure 11

### Supplementary Figure 11

| Gene Promoter | Forward and Reverse primer sequences |
|---|---|
| APOBEC3A | 5'-AGAGTGGGCACATCAAAACC-3'<br>5'-AACCTGGAAGCTGCTGAGTC-3' |
| APOBEC3B | 5'-GGTCACTTTAAGGAGGGCTGT-3'<br>5'-TAGATACGCTTGTCCCTGTCC-3' |
| APOBEC3C | 5'-AACCAGAAAGAGGGCCAGAG-3'<br>5'-TTACAGCGTCCTTGCAGTTG-3' |
| APOBEC3D | 5'-CCTACACCAGCGCCTGAG-3'<br>5'-GTGAGAGAGCGAGGCTTCC-3' |
| APOBEC3F | 5'-CAGACACCTGGCCCTTTACT-3'<br>5'-CCTCCTCTCCACCATCAAGA-3' |
| APOBEC3G | 5'-AGACGCCTGGCCATTTACT-3'<br>5'-CCTCCTCTCCACCATCAAGA-3' |
| APOBEC3H | 5'-CTCCAGTCCCACAAAAGGAA-3'<br>5'-TTCACTTTTGGCAGCTCTCC-3' |
| mCdkn1a (p21) | 5'-GAGACCAGCAGCAAAATC -3'<br>5'-CAGCCCCACCTCTTCAATTC-3' |
| mMdm2 | 5'-TGACTCAGCTCTTCCTGTGG-3',<br>5'-GAGCTGTCCCTTACCTGGAG-3' |

**Supplementary Figure 11: ChIP qPCR primers for the 7 hA3 genes and mouse *Cdkn1a* and *Mdm2.***
